# Natural Occurring Non-Synonymous Single Nucleotide Polymorphisms in Integrase and RNase H Regulate Assembly and Autoprocessing of HIV-1

**DOI:** 10.1101/2021.03.15.435559

**Authors:** Tomozumi Imamichi, John G. Bernbaum, Sylvain Laverdure, Jun Yang, Qian Chen, Helene Highbarger, Ming Hao, Hongyan Sui, Robin Dewar, Weizhong Chang, H. Clifford Lane

**Affiliations:** Laboratory of Human Retrovirology and Immunoinformatics, Applied and Developmental Directorate, Frederick National Laboratory, Frederick, Maryland, 21702, USA; Integrated Research Facility, National Institute of Allergy and Infectious Diseases, National Institutes of Health, Frederick, Maryland, USA; Virus Isolation and Serology Laboratory, Applied and Developmental Directorate, Frederick National Laboratory, Frederick, Maryland, USA; Laboratory of Immunoregulation, National Institute of Allergy and Infectious Diseases, National Institutes of Health, Bethesda, Maryland, USA

## Abstract

Recently, a genome-wide association study using plasma HIV RNA reported that 14 naturally occurring non-synonymous single nucleotide polymorphisms (SNPs) in HIV derived from anti-retrovirus naïve patients were associated with virus load (VL). However, the impact of each mutation on viral fitness was not investigated. Here, we constructed a series of HIV variants encoding each SNP using site-directed mutagenesis and examined their replicative abilities and biological properties. An HIV variant containing Met-to-Ile change at codon 50 in integrase (HIV(IN:M50I)) was found an impaired virus. Despite the mutation being in integrase, a quantification assay demonstrated that the virus release was significantly suppressed (P<0.001). Transmission electron microscopy analyses revealed that the accumulation of abnormal shapes of buds on the plasma membrane and the released virus particles retained immature forms. Western blot analysis demonstrated a defect in autoprocessing of GagPol and Gag polyproteins in the HIV(IN:M50I) particles. Förster Resonance Energy Transfer (FRET) assay displayed that GagPol containing IN:M50I (GagPol(IN:M50I)) significantly increased the efficiency of homodimerization (P<0.05) and heterodimerization with Gag (P<0.001), compared to GagPol(WT). HIV replication assay using a series of variants of HIV(IN:M50I) elucidated that the C-terminus residues, Asn at codon 288, plays a key role in the defect and the impaired maturation and replication capability was rescued by two other VL-associated SNPs, Ser-to-Asn change at codon 17 in integrase or Asn-to-Ser change at codon 79 in RNase H. These data demonstrate that Gag and GagPol assembly, virus release and autoprocessing are not only regulated by integrase but also RNase H.

**Importance:** A nascent HIV-1 is noninfectious. To become an infectious virus, Gag and GagPol polyproteins in the particles need to be cleaved by mature HIV protease (PR). PR is initially translated as an inactive embedded enzyme in a GagPol polyprotein. The embedded PR in homodimerized GagPol polyproteins catalyzes a proteolytic reaction to release the mature PR. This excision step by a self-cleavage is called autoprocessing. Here, during the evaluation of roles of naturally emerging non-synonymous SNPs in HIV RNA, we found that autoprocessing is inhibited by Met-to-Ile change at codon 50 in integrase in GagPol which increases the efficiency of heterodimerization with Gag. This defect was recovered by co-existing of other SNPs: Ser-to-Asn change at codon 17 in integrase or Asn-to-Ser mutation at codon 79 in RNase H, suggesting that autoprocessing is regulated by not only integrase but also RNase H in GagPol polyprotein.

## Introduction

Nascent human immunodeficiency virus type 1 (HIV-1) particles are released from infected host cells as immature and non-infectious viruses (1). These immature viral particles contain Gag and GagPol polyproteins, accessory proteins including Nef precursor, and genomic viral RNAs. The Gag and GagPol polyproteins are composed of viral structural proteins and the viral enzymes: protease (PR), reverse transcriptase (RT), RNase H (RH), and integrase (IN). During proteolytic maturation, the polyproteins in the immature particle are cleaved by the viral PR at a total of 11 sites on the Gag and GagPol polyproteins and one site on the Nef precursor (1–3), and then the immature particle is converted into its infectious mature form. While these catalytic activities of PR are well-described, there are other aspects of PR that may play a role in viral replication. During viral Gag and GagPol assembly and budding at the cell membrane, GagPol polyproteins dimerize, followed by dimerization of the PR domains of the GagPol dimer, leading to the release of a functional dimerized PR in the viral particles. Inhibition of dimerization of the GagPol polyproteins has been considered a unique therapeutic target (4). Autoprocessing, that is, the initial activity of releasing the embedded PR in the GagPol polyprotein by its own processing results in the formation of mature PR (5–7) that can catalyze the other cleavage reactions. Therefore, the homodimerization of the GagPol polyproteins in the immature virion is required for this excision of PR. IN is a component of the GagPol polyprotein and a biologic role of the integrase protein of HIV-1 has been thought to be limited to inducing integration of proviral DNA to the host cell genome following infection. However, *in vitro* studies have demonstrated that drug resistance mutations to the experimental integrase inhibitor, KF116, interfere with autoprocessing and thus have suggested a potential role for IN in autoprocessing of GagPol polyprotein (8, 9).

In a recent sub-study of the Strategic Timing of Antiretroviral Therapy (START) study (10), the investigators identified a series of 14 naturally-occurring non-synonymous single nucleotide polymorphisms (SNPs) in HIV-1 that correlated with different levels of viremia in treatment-naïve patients (11). However, the specific impact of each amino acid (aa) substitution on viral fitness was not investigated. In the present study, we created a series of recombinant viruses containing each of the SNPs using site-directed mutagenesis and characterized the role of each mutation in viral fitness. Of all the mutants, only the Met-to-Ile substitution at codon 50 of IN (IN:M50I) resulted in the loss of replication capability leading to the release of abnormally shaped virions and interfering with autoprocessing. Of note, this “off-target” effect was restored by compensatory mutations in IN(IN:S17N) or RH(RH:N79S) in the 14 SNPs. These results identify critical roles for RH and IN in the autoprocessing of the HIV-1 GagPol polyprotein.

## Results

### Impact of SNPs on Virus Replication Fitness

A total of 14 non-synonymous SNPs were found to correlate with plasma viral loads in a cohort of 3,592 anti-retroviral drug-naïve HIV-infected patients (11). To characterize the impact of each aa substitution (Table 1) on viral fitness, we constructed a series of HIV variants containing each change using site-directed mutagenesis using the cloned HIV laboratory strain, HIV_NL4.3_. Since this strain contains an Asn-to-Ser substitution at codon 79 of RH (RH:N79S), this site was back-mutated from Ser to Asn to yield a clone equivalent to the RH of the clade B consensus sequence (RH:N79). We used this construct, pNL(WT), as the backbone wild-type HIV (HIV(WT)) control in the current study. Each non-synonymous SNP was introduced onto the WT control backbone using site-directed mutagenesis.

**Table 1:**
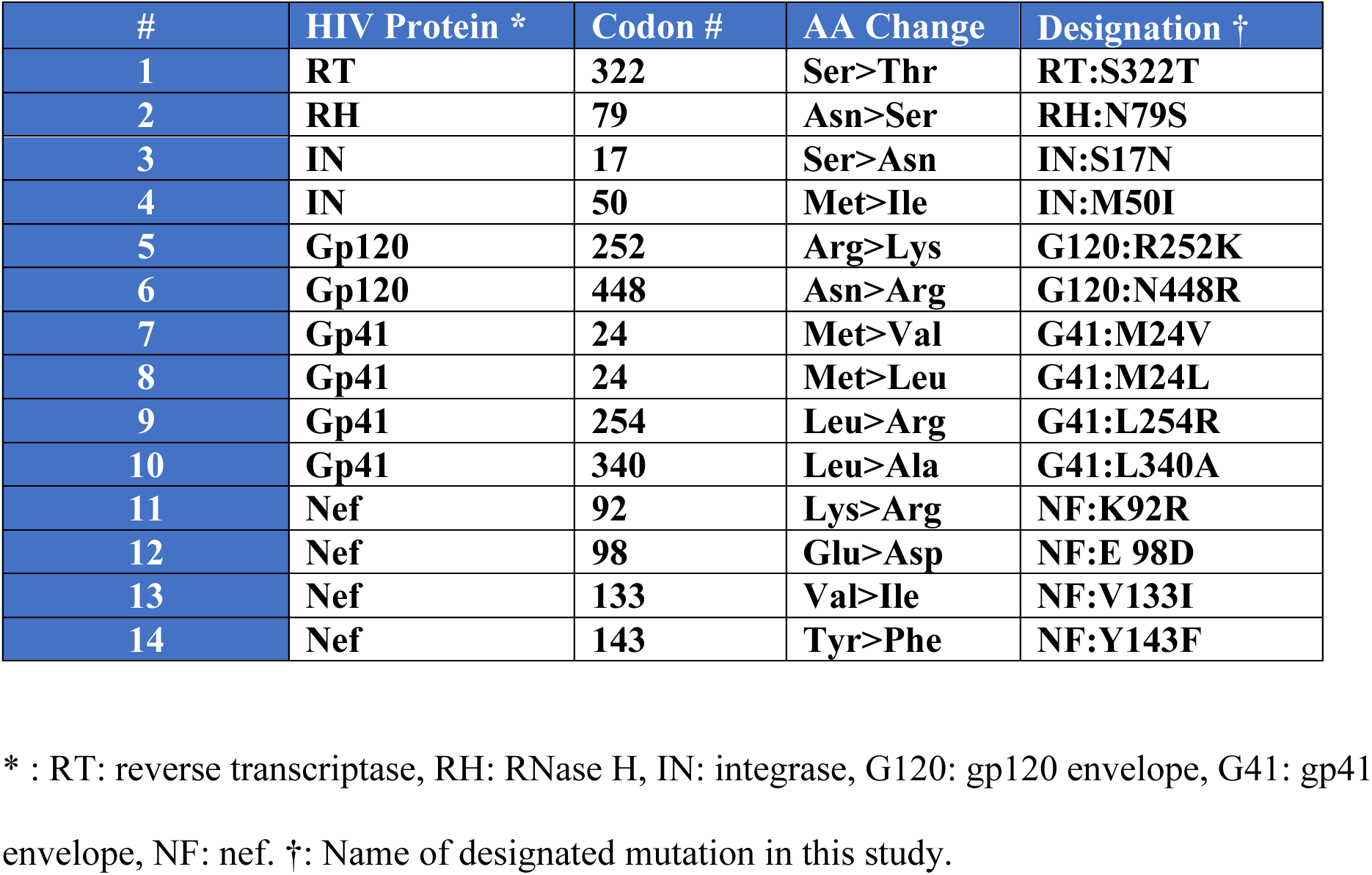
Natural Polymorphic Mutations Associated with Viral Loads in Antiviral drug naïve HIV-infected patients.

Plasmids encoding each SNP were transfected into HEK293T cells, and then viral stocks were prepared as described in the Materials and Methods. Pelleted viral particles were resuspended in 1/100 volume of the starting materials and used as viral stocks. To determine the amounts of virus in each variant stock, HIV p24 antigen concentrations in each stock were measured. Almost all variants except HIV(IN:M50I) demonstrated comparable amounts of p24 concentrations with that of HIV(WT) (89.0 ± 12 μg/ml, *n*=7) (Supplementary Table S2). In striking contrast, the p24 concentrations of HIV(IN:M50I) stocks were 0.30 ± 0.087 μg/ml (*n*=7) and thus less than 0.3 % of that in HIV(WT) (P<0.001). To define whether the suppression of HIV(IN:M50I) release is caused by only HEK293T cells, Hela cells were also transfected with HIV(WT) or HIV(IN:M50I) construct and then released HIV amounts were quantified. HIV(IN:M50I) virus was released by 0.3 % of HIV(WT) (Supplementary Table S2) from the cells, indicating that the inhibition is not caused in HEK293T specific manner. It is known that Hela cells produce Interferons (IFNs) in response to transfection plasmid DNA as an innate immune response, while HEK293T cells lack the inducting activity (12). To avoid any impact of IFNs in downstream experiments, we used HEK293T cells in entire studies as an HIV-producing cell.

To investigate the impact of each mutation on viral replication, we infected PHA-stimulated primary CD4+ T cells from three independent donors with each HIV variant, and viral replication was monitored for 14 days. All mutants except HIV(IN:M50I) replicated comparably to HIV(WT), and the HIV(IN:M50I) variant showed little to no replication (Fig. 1a – 1c, Supplementary Fig. S1a – S1d). Compared to HIV(WT), in the presence of IN:M50I mutation, HIV replication was suppressed by 99.9 ± 0.030 % (n=7, P<0.01) on the 7^th^ day after infection, and even when the HIV(IN:M50I)-infected cells were cultured for additional 7 days, significant viral replication was not detected.

**Figure 1.**
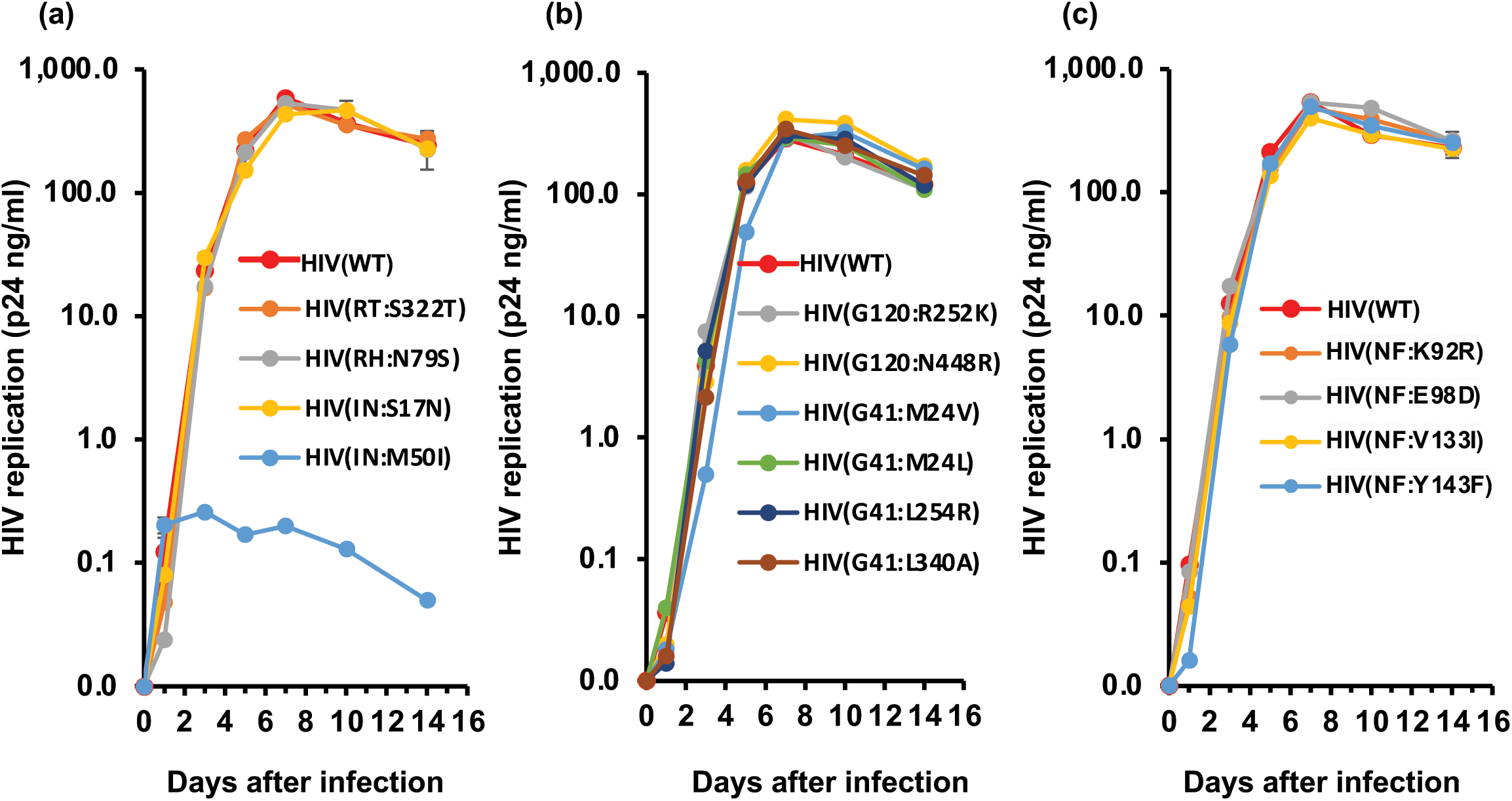
Characterization of 14 SNPs on HIV replication. (A-C) PHA-stimulated primary CD4(+) T cells from normal healthy donor were infected with 10 ng p24 amounts of HIV(WT) or variants containing mutation in GagPol (A), envelope (B), or Nef (C) as described in the materials and methods. The infected cells were cultured for 14 days with media changed every 3-4 days. HIV replication was monitored using a p24 antigen capture kit. Representative data from three independent assays are presented as mean ± SD (n=3).

To further investigate the biologic properties of the IN:M50I mutant, we examined the morphology of the transfected HEK293T cells and virus particles 24 hours after transfection using transmission electron microscopy (TEM). As shown in Fig. 2a and 2b, both HIV(WT) and HIV(IN:M50I)-transfected cells led to the release of viral particles. Viral particles from the HIV(IN:M50I) transfected cells were larger and more variable in size than particles from wild-type transfected cells. These differences persisted after an additional 24 hours of culture and further accumulation of virus buds on the cell surface was observed in HIV(IN:M50I) (Supplementary Fig. S2a and S2b).

**Figure 2.**
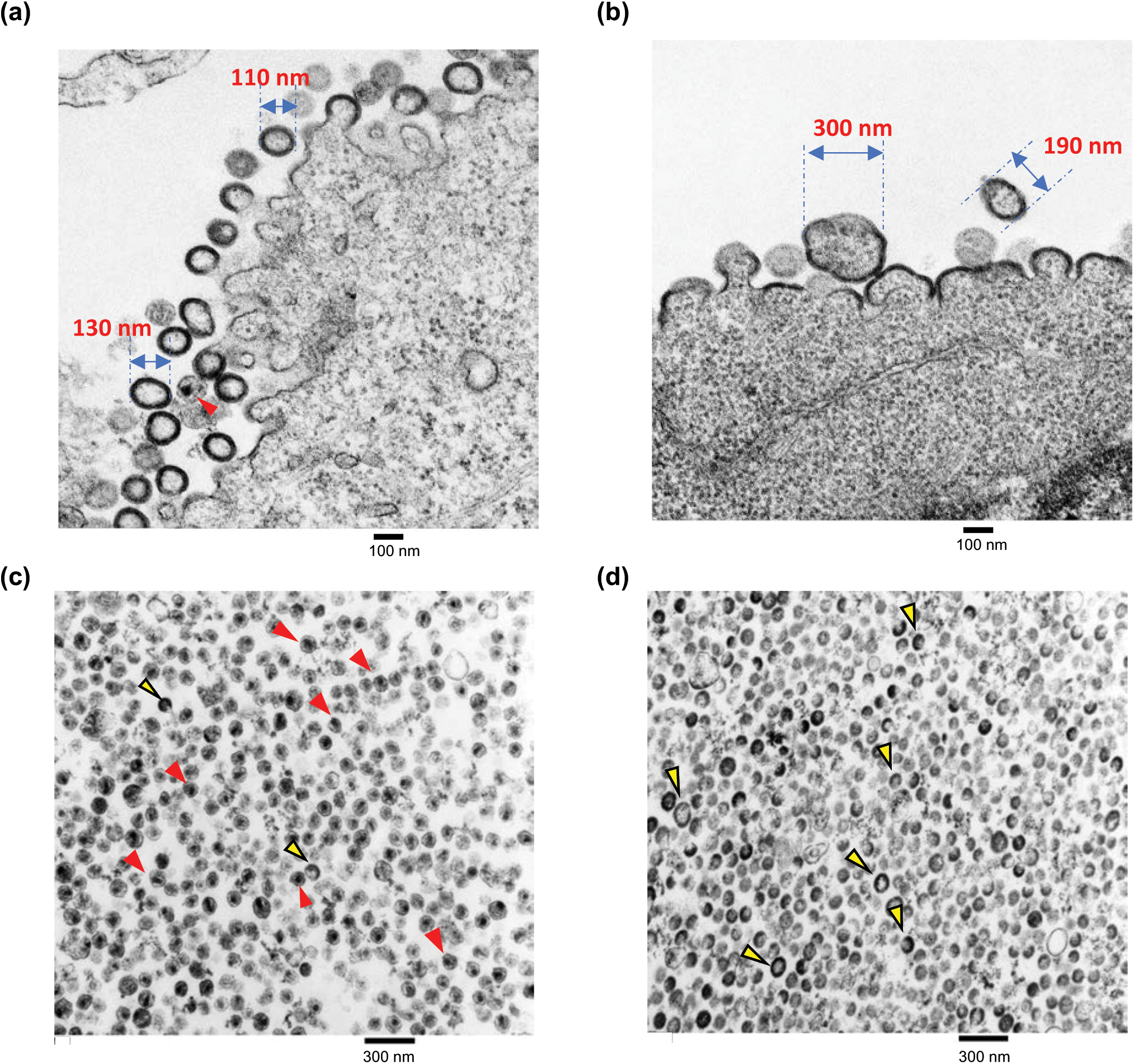
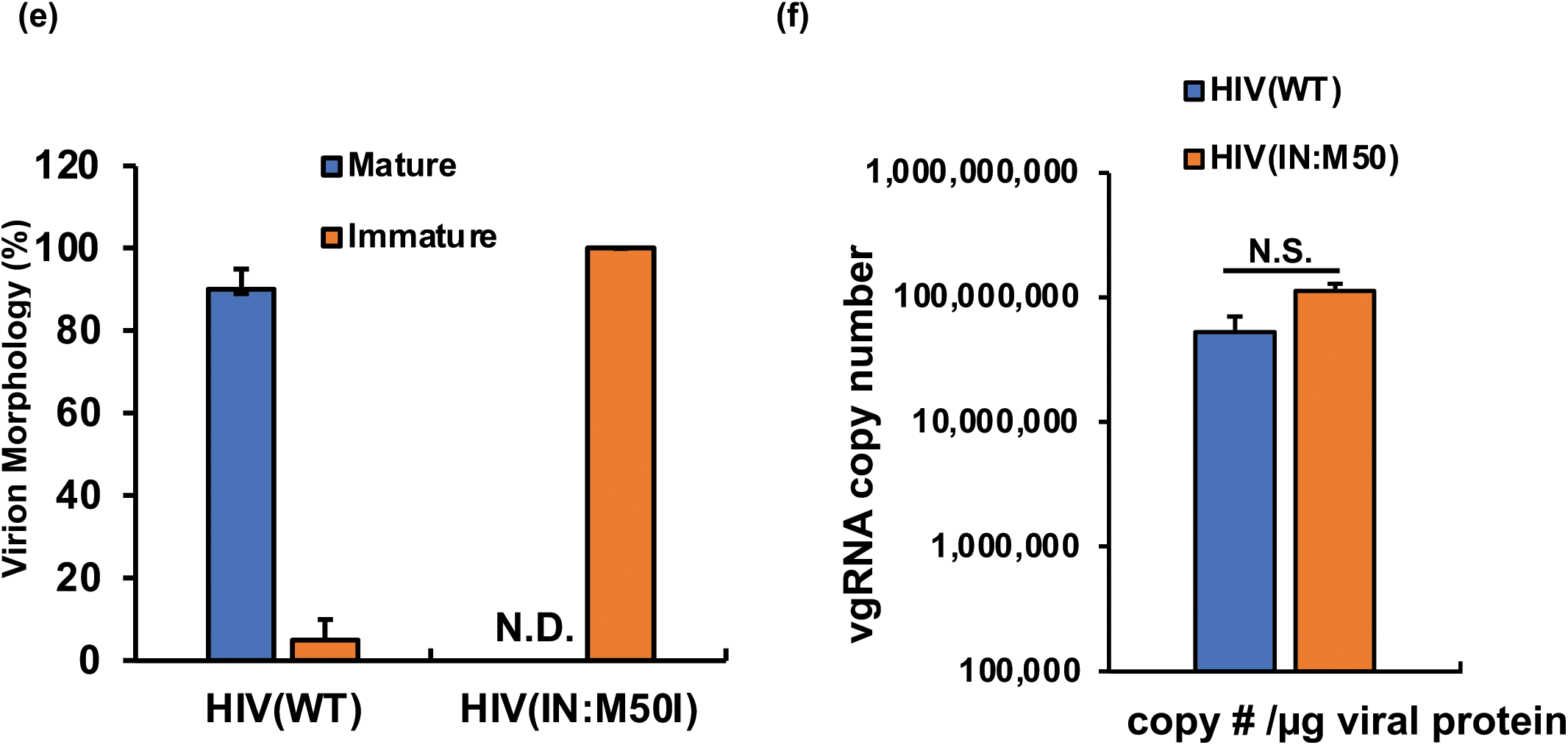
Characterization of viral morphology. HEK293T cells were transfected with plasmids encoding HIV(WT) (A) or HIV(IN:M50I) (B) and cultured for 24 hours. Cells were fixed and then TEM images were taken as described in the Materials and Methods. (c and d) TEM image analyses of sucrose-purified HIV particles of HIV(WT) (C) and HIV(IN:M50I) (D). Red and green arrows heads indicate typical electron-dense particles with a conical core (red) and electron-dense particles without a core (green). Scale bars show 100 nm in (A) and (B), 200 nm in (C) and (D). (E) Relative population of viral morphology was calculated from 1000 particle images by three independent assays. Data indicates mean ± SE (n=3). (F) Viral genomic RNA copy numbers in viral stocks were compared between HIV(WT) and HIV(IN:M50I). Copy numbers were determined using the RealTime HIV-I assay kit from three independent virus stocks, and data were normalized to total viral protein. Data shows mean ± SE (n=3).

TEM of HIV(WT) particles revealed virions with the expected outer envelope; 90-95 % having an electron-dense core (Fig. 2c and 2e). In contrast, TEM of HIV(IN:M50I) particles revealed a similar outer envelope; with no evidence of a core (Fig. 2d and 2e). In addition, while the WT particles were relatively uniform in size (110 – 130 nm in diameter); the M50I particles were highly variable in size (190 – 300 nm in diameter). Taken together these findings indicate the HIV(IN:M50I) variant exhibited abnormalities in both virion formation and unlike HIV(WT) with its dense core, the mutant viruses had ring, doughnut-shaped or teardrop structures, suggesting an irregular assembly of viral proteins and lack of PR activity in the HIV(IN:M50I) virus particles.

Gag and GagPol polyproteins play an important role in the intra-cytoplasmic trafficking of viral genomic RNAs (vgRNAs) and their eventual incorporation into the immature budding virions (1, 13–15). To determine whether or not the mutant phenotype of IN:M50I virions was due to a defect in the delivery of vgRNAs to the budding viral particles, we measured vgRNA copy number using purified HIV(WT) and HIV(IN:M50I) particles. No significant differences were observed in vgRNA copy number per microgram of virion protein between HIV(WT) and HIV(IN:M50I) (P=0.066, n=3) (Fig. 2f), indicating that the IN:M50I mutation had no impact on the incorporation of vgRNAs into virions.

Given the immature appearance of IN:M50I virions, we next sought to determine whether the mutation led to any changes in autoprocessing and proteolytic cleavage of the GagPol polyproteins. Purified particles were analyzed for viral proteins by Western blot (WB) using a series of antibodies specific to p24 capsid protein (CA), PR, RT, IN or Nef. In contrast to the wild-type virions, HIV(IN:M50I) virions contained uncleaved Gag and GagPol polyproteins and immature Nef and did not contain the cleaved fragments of CA, PR, RT, and IN as detected in HIV(WT) (Fig. 3a – 3e). These results indicated that the IN:M50I mutation was somehow interfering with the activity of the HIV-1 PR, likely at the autoprocessing step.

**Figure 3.**
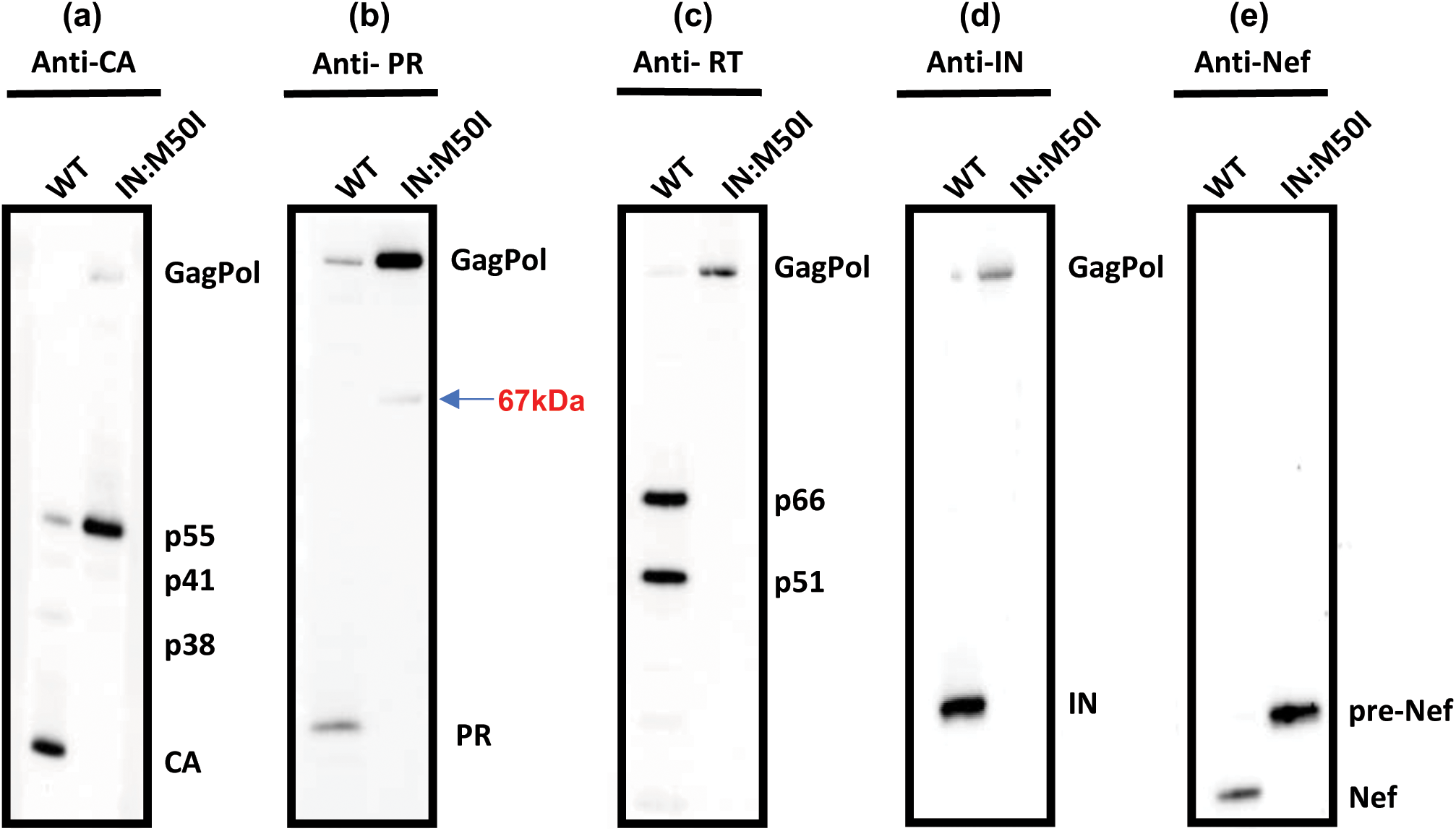
Evaluation of Gag ang GagPol processing. A total of 1μg of viral proteins from HIV(WT) and HIV(IN:M50I) particles were subjected for WB using antibodies to CA (p24), PR, RT, IN or Nef. Protein bands were detected by using the ECL assay as described in the Materials and Methods. The protein gel for detecting PR was run in MES running buffer (Thermo Fisher), other gels were run in MOPS running buffer (Thermo Fisher). Data are representative of two independent experiments.

### Impact of IN:M50I mutation on GagPol dimerization

The autoprocessing of GagPol polyprotein is dependent upon the formation of GagPol polyprotein homodimers (5, 16), thus we hypothesized that an IN:M50I mutation might lead to a structural hindrance in GagPol interfering with homodimerization. To determine whether or not the M50I mutation suppresses homodimer formation, we employed a Förster resonance energy transfer (FRET) assay using the plasmids pGag(MA/EGFP/CA), pGagPol(MA/mSB/CA), pGag(MA/mSB/CA), pGagPol(MA/EGFP/CA) (17). Since the pGag(MA/EGFP/CA) and pGagPol(MA/mSB/CA) contained RH:N79S, they were back-mutated to RH:S79N, and IN:M50I was induced in both plasmids.

We first assessed the efficiency of homodimerization among Gag-to-Gag (G-G), GagPol(WT)-to-GagPol(WT) and GagPol(IN:M50I)-to-GagPol(IN:M50I). Consistent with another report(17), FRET signals from G-G (Fig. 4a) were stronger than the signals from other pairs (Fig. 4b and 4c) and predominantly located on the cell membranes. The efficiency of the homodimerization of GagPol(WT)-to-GagPol(WT) and the GagPol(IN:M50I)-to-GagPol(IN:M50I) were 31 ± 3.4 % (n=3, P<0.001) and 49 ± 3.0 % (n=3, P<0.001) of that of the Gag homodimerization, respectively (Fig. 4f). Interestingly, GagPol(IN:M50I) homodimerization had a 1.7 ± 0.28-fold higher efficiency than did GagPol(WT) homodimerization (n=3, P<0.05). Next, we compared the efficiency of heterodimerization between GagPol(WT) and Gag and between GagPol(IN:M50I) and Gag. GagPol(WT) formed a heterodimer with Gag on the plasma membrane with 36 ± 2.8 % (n=3, P<0.001) of the efficiency of the Gag homodimerization (Fig. 4a, 4d and 4g) and the heterodimer localization at the cytoplasm was observed. Surprisingly, the efficiency of the GagPol(M50I)-to-Gag heterodimerization was indistinguishable (89 ± 5.0%) from that of Gag homodimerization (n=3, P=0.17) (Fig. 4a, 4e and 4g), and the efficiency of the heterodimerization of GagPol(IN:M50I)-to-Gag was 2.5 ± 0.10-fold higher than that of GagPol(WT)-to-Gag (n=3, P<0.001) on the membrane, suggesting that in the presence of IN:M50I mutation, GagPol may preferentially form a heterodimer with Gag on the plasma membrane. To clearly compare the distribution of Gag, GagPol(WT) and GagPol(IN:M50) in cells, we transfected each GFP-construct into HEK293T cells and analyzed protein distribution. Gag and GagPol(IN:M50) were distributed throughout cells (the cytosol and the plasma membrane) (Supplementary Fig S3a and S3c), while GagPol(WT) was predominantly located in the cytosol with an aggregated form (Supplementary Fig S3b), indicating that GagPol(IN:M50I) can distribute on the plasma membrane by itself, while GagPol(WT) by itself mainly localizes in the cytosol.

**Figure 4.**
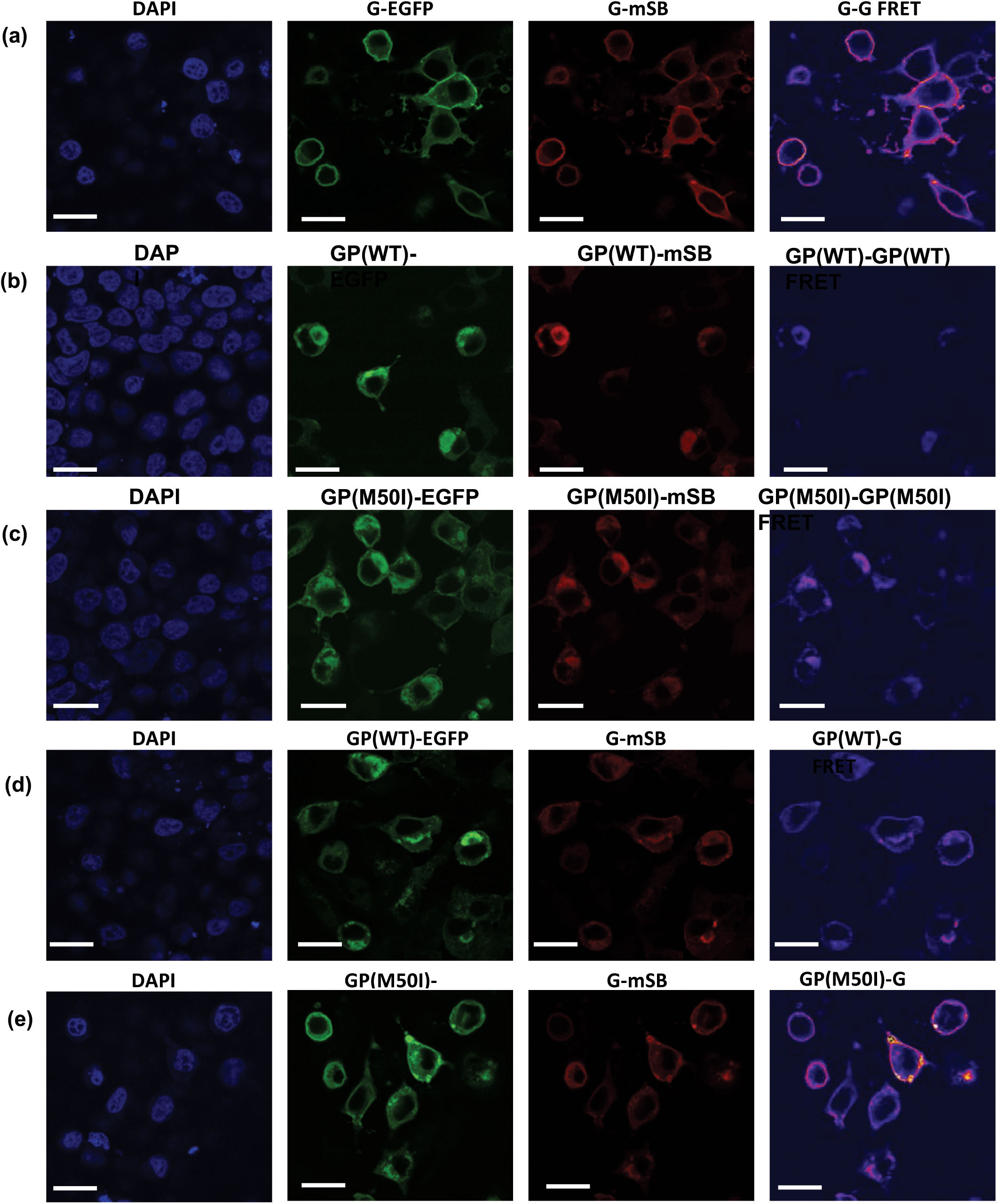
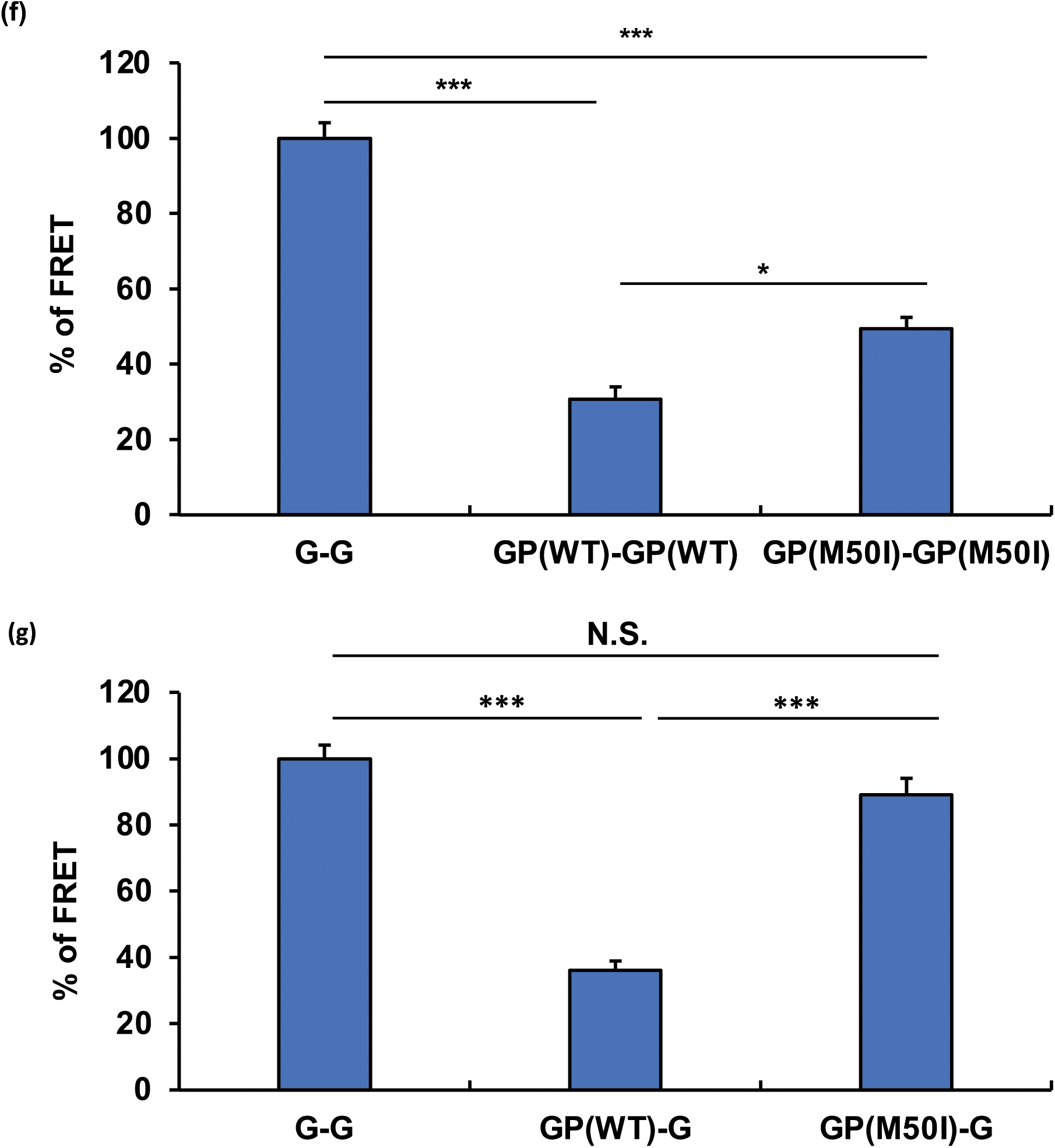
Evaluation of Gag and GagPol dimerization. HEK293T cells were co-transfected with pGag-GFP and pGag-mSB (A), pGagPol(WT)-GFP and pGagPol(WT)-mSB (B), or pGagPol(IN:M50I)-GFP and pGagPol (IN:M50I)-mSB (C), pGagPol(WT)-GFP and pGag-mSB (D) or pGagPol(IN:M50I)-GFP and pGag-mSB (E), and then FRET assays were conducted as described in the Materials and Methods. In the figures, Gag-GFP, Gag-mSB, GagPol(WT)-GFP, GagPol(WT)-mSB, GagPol(IN:M50I)-GFP, GagPol(IN:M50I)-mSB are displayed G-GFP, G-mSB, GP(WT)-GFP, GP(WT)-mSB, GP(M50I)-GFP and GP(M50I)-mSB, respectively. (F and G) FRET efficiencies calculated as described in the Materials and Methods. Gag or GagPol homodimerization are described G/G and GP/GP. Gag and GagPol heterodimerization is depicted G/GP. The efficiencies of GP/GP (F) and G/GP combinations (G) were compared with that of the G/G pair. Data indicate mean ± SE from five independent assays. * P<0.05, *** P<0.001.

Given that GagPol(IN:M50I) can form a homodimer, it was considered that the inhibition of autoprocessing may be caused by not only the higher efficiency of formation of the heterodimer but also other factors such as a change in the binding affinity of embedded PR in GagPol to the initial cleavage site in GagPol polyproteins. In the WB result using anti-PR antibody (Fig. 3b), an ∼67 kDa polypeptide band was consistently detected in HIV(IN:M50I) virions as an additional unique cleaved product. HIV PR cleaves a total of 9 cleavage sites in GagPol polyprotein (Fig. 5a) (1). It is reported that HIV PR cleaves GagPol in an ordered sequence leading to virus maturation (18). Generally, it is considered that the initial cleavage occurs between p2 spacer peptide and the NC protein (at cleavage site 3 in Fig. 5a) in HIV(WT) (19–25) as an intramolecular reaction (*cis*-reaction) (5, 16, 26, 27). Since the unique ∼67 kDa polypeptide band was detected by anti-PR antibody, we presumed that the band may result from the cleaved product between PR and RT at cleavage site 7 (Fig. 5a) due to an abnormal cleavage order and that the ∼67 kDa polypeptide may be composed of MA/CA/p2/NC/TF/p6*/PR (Fig. 5a), implicating that the IN:M50I mutation causes a disorder in the cleavage sequence of autoprocessing of GagPol. To address this hypothesis, we created mutant viruses in which cleavage site 7 in HIV(WT) and HIV(IN:M50I) was changed from Phe-Pro to Val-Pro by point mutagenesis and the resulting viruses were designated HIV(WT_Δ7) and HIV(M50I_ Δ7), respectively. It is reported that virus lacking cleavage site 7, which makes a fusion protein of PR and RT (PR-RT, Fig.5a), still possesses a functional PR activity (28). We therefore expected that if the IN:M50I mutation had no direct impact on PR activity, HIV(M50I_Δ7) would function as well as HIV(WT_Δ7) at GagPol processing. Those constructs were transfected into HEK293T cells, viral particles were collected, and then WB analysis was performed using those virus lysates with a polyclonal anti-PR antibody. As expected, the 67 kDa band was no longer present in HIV(M50I_Δ7) (Fig. 5b), instead dominant bands at 73 kDa and 90 kDa with minor bands at 16 and 11kDa were detected, which were also detected at a comparable level in HIV(WT_Δ7). WB analysis using anti-CA antibody and anti-IN antibody were also conducted, and comparable levels of mature CA- and IN-sized bands were detected in both HIV (WT_Δ7) and HIV (IN:M50I_ΔD7) (Fig. 5c and 5d). These findings indicated that the IN:M50I mutation alters the order of the autoprocessing rather than directly inhibiting PR function and consequently maturation of the released virions fails due to inhibition of the initial cleavage at cleavage site 3.

**Figure 5.**
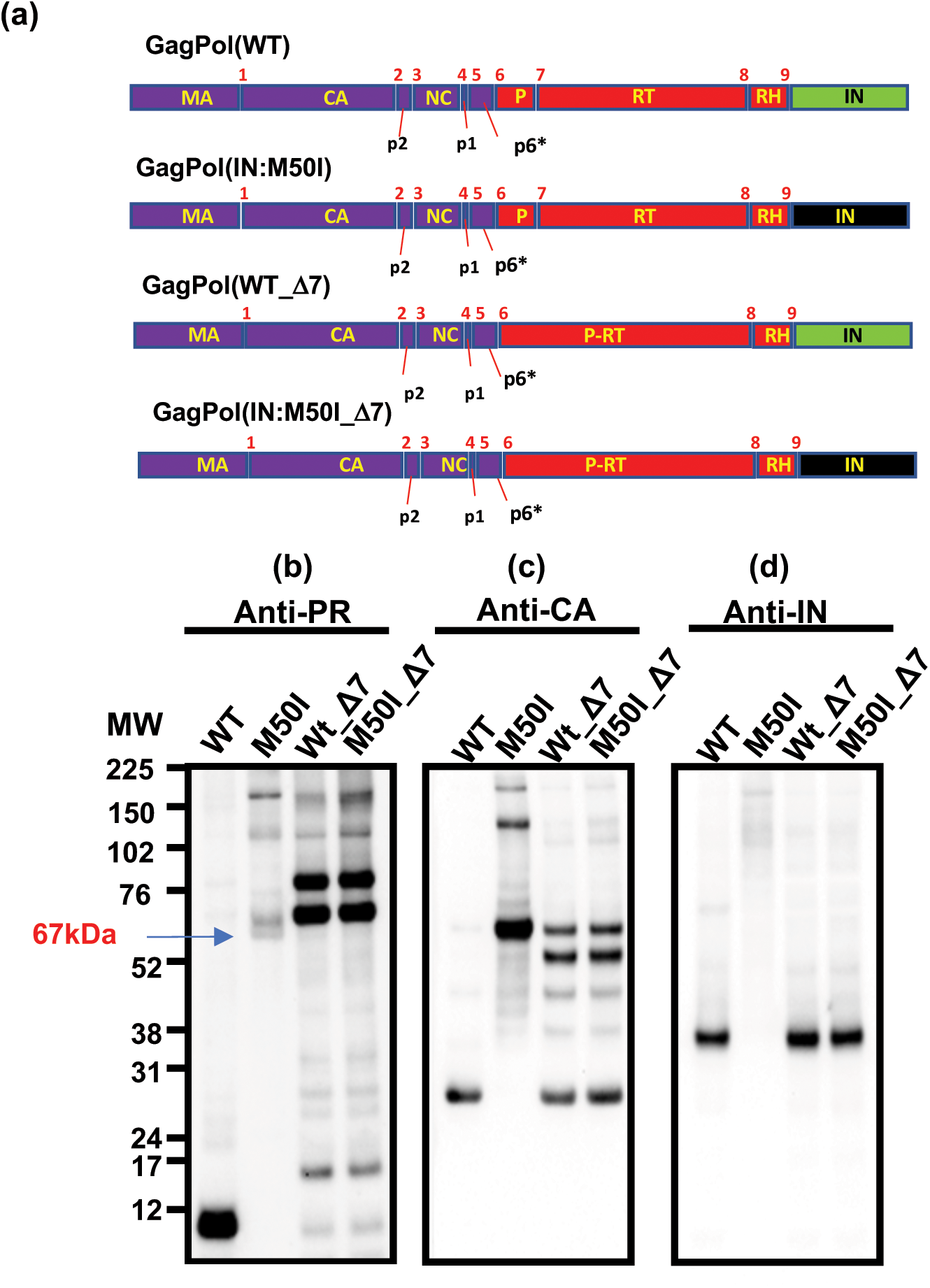
Effect of the deletion of the cleavage site between PR and RT in GagPol on autoprocessing. (A) Diagrams indicate structure of GagPol (WT) and variants. Numbers above the diagrams indicate cleavage sites #1 – #9 by mature PR. The blue dot square demonstrates a predicted fragment of 67 kDa detected in HIV(IN:M50I) by anti-PR antibody. Molecular size of each domains: MA=17 kDa, CA=24 kDa, p2=2 kDa, NC=7 kDa, TF=1 kDa, P=10 kDa, RT=51 kDa, RH=10 kDa, IN=35 kDa. The cleavage site 7 between PR and RT in HIV(WT) and HIV(IN:M50I) was deleted by a point mutagenesis as described in the Materials and Methods and the resulted clones contain a fusion gene of PR and RT (PR-RT). (B-D) Virus particles are isolated using ultracentrifugation as described in the Materials and Methods, and viral lysates were subjected for WB. PR, CA and IN were detected by anti-PR, anti-HIVp24(CA), and anti-IN antibody, respectively.

### Identification of Compensatory Mutations

Our viral fitness results demonstrated that the IN:M50I mutation was a lethal mutation when introduced as a single change, however, since it was identified from a study of circulating virions in anti-retroviral drug-treatment-naïve patients, we postulated that the circulating viruses must also contain a compensatory mutation(s) elsewhere in the genome. It has been reported that HIV_NL4.3_ carrying IN:M50I mutation is replication-competent *in vitro* (29) and HIV_NL4.3_ exhibits RH:N79S polymorphism (30). This variant was also found among the quasi-species from the patient pool from which IN:M50I was identified (Table1). Thus, we presumed that RH:N79S may be a compensatory mutation for IN:M50I. Also, among the swarm of replicating viruses, another mutation in IN, IN:S17N, was also present, suggesting that this might also be a compensatory mutation. To test these hypotheses, we constructed variants carrying a combination of IN:M50I and RH:N79S (HIV(RH:N79S/IN:M50I)), or IN:M50I and IN:S17N (HIV(IN:S17N/IN:M50I)) and analyzed their ability to replicate. Both double-mutant virions restored the ability to replicate (Fig. 6a and 6b, Supplementary Fig. S4a and S4b) and led to the formation of mature virions containing mature PR and CA proteins (Fig. 6c and 6d) in WB and amounts of released virus from the producing cells were comparable to that of the wild-type (Supplementary Table S2). TEM demonstrated that the viral particles of the double mutants were indistinguishable from that of HIV(WT) (Fig. 6e and 6f).

**Figure 6.**
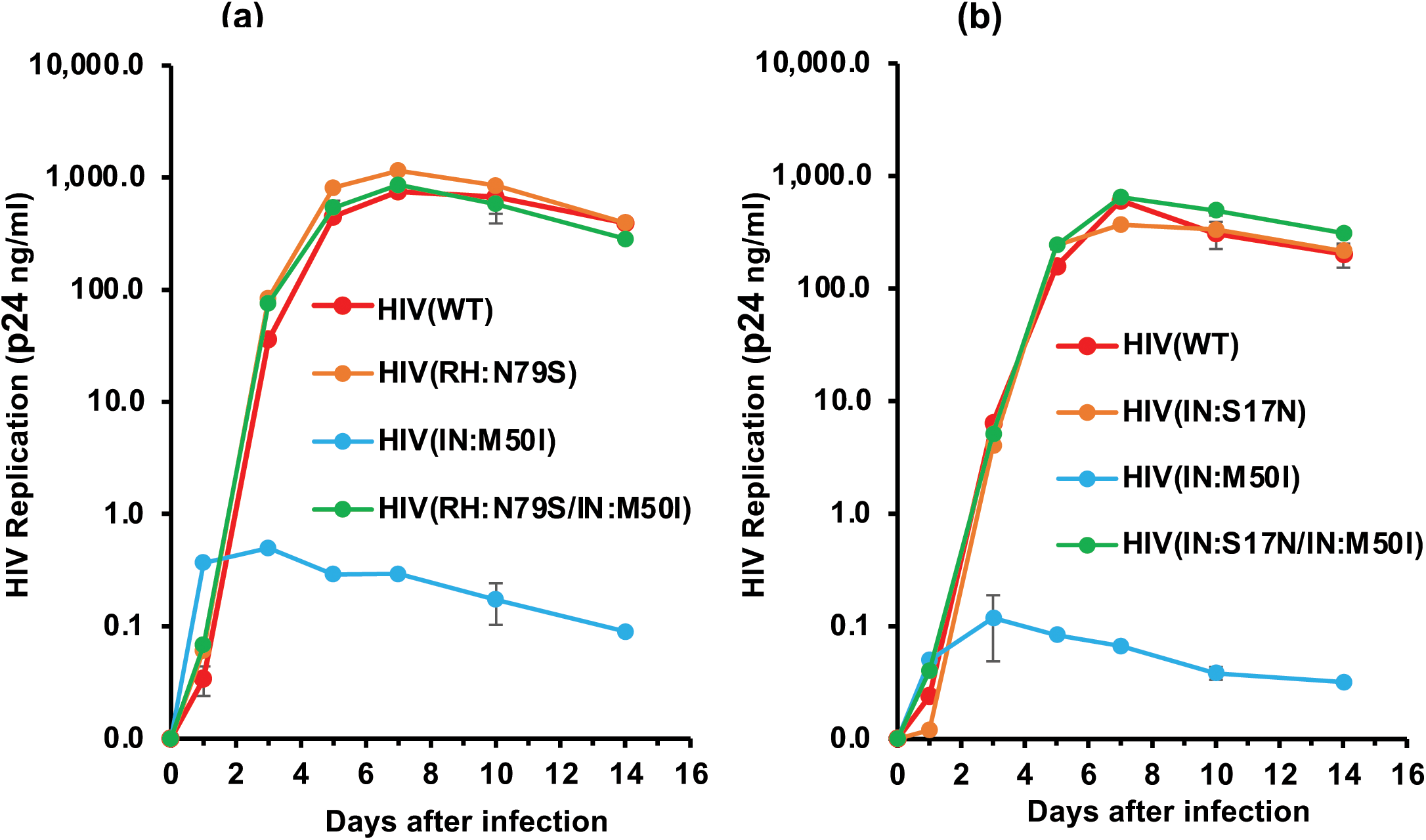
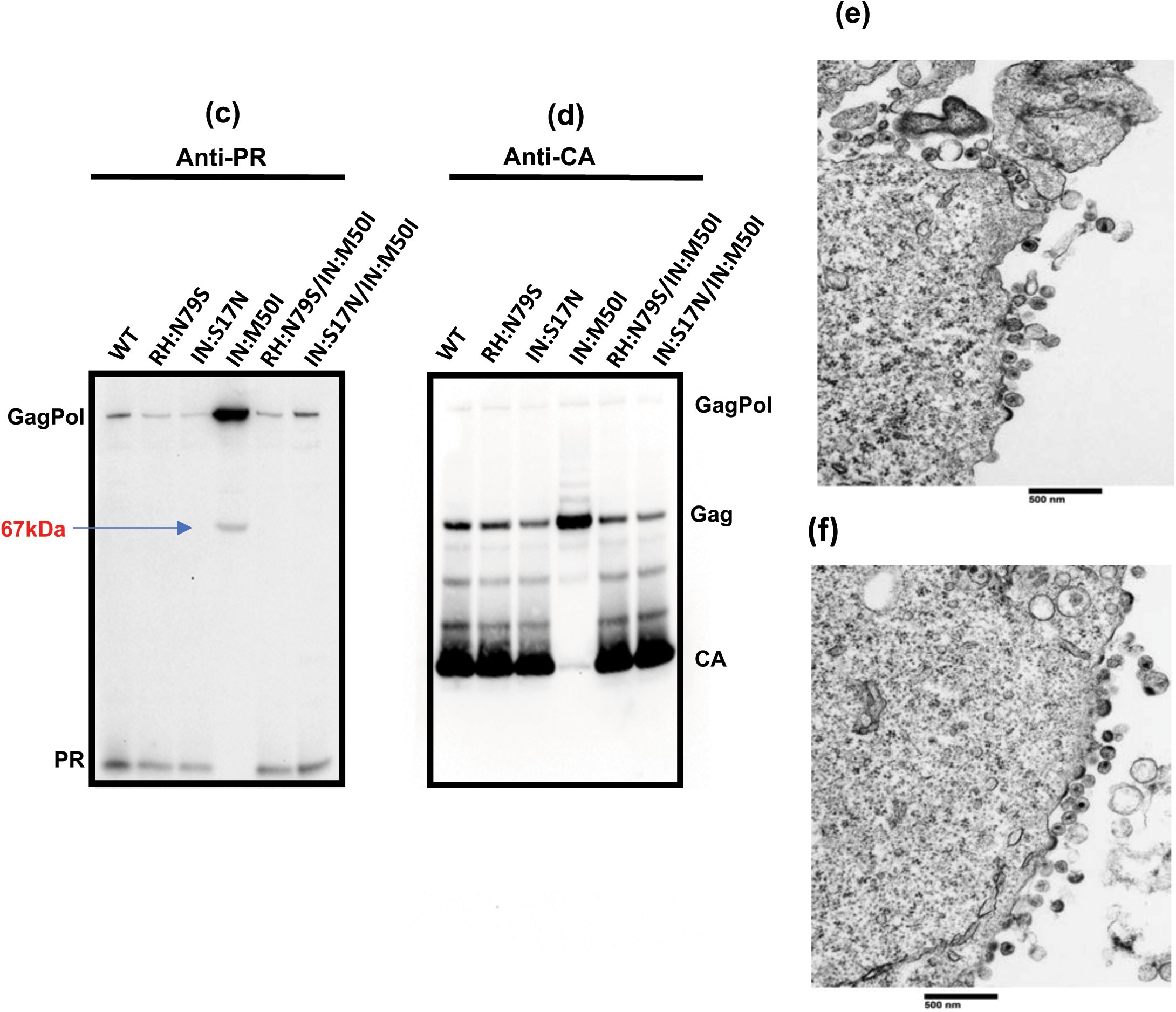
Evaluation of RH:N79S or IN:S17N mutation on HIV(IN:M50I) virus replication and GagPol processing. (A and B) PHA-stimulated primary CD4(+) T cells were infected with HIV(WT), HIV(IN:M50I), HIV(RH:N79S), or HIV(RH:N79S/IN:M50I) (A) or with HIV(IN:S17N) or HIV(IN:S17N/IN:M50I) (B). The infected cells were cultured for 14 days with media changed every 3-4 days. HIV replication was monitored using a p24 antigen capture kit. Representative data from two independent assays are presented. Data are presented as mean ± SD (*n*=3). (C and D) Viral lysates of HIV(WT), HIV(RH:N79S), HIV(IN:S17N), HIV(IN:M50I), HIV(RH:N79S/IN:M50I) and HIV(IN:S17N/IN:M50I) underwent WB analyses using anti-PR (C) and anti-CA antibodies (D). (E and F) Comparison of morphology of viral particles using TEM. HEK293T cells were transfected with plasmid DNA encoding HIV(RH:N79S/IN:M50I) (E) or HIV(IN:S17N/M50I) (F) and then cultured for 24 hours. Cells were fixed for TEM analysis.

To determine the potential clinical relevance of these findings, we analyzed the population of viruses carrying these mutations using the Los Alamos HIV Sequence Database (31). Of a total of 5100 HIV sequences, 401 (7.9 % of the total sequences), carried the IN:M50I mutation (Supplementary Table S3 and S4). Among these 401 sequences, 252 sequences carried either the RN:N79S or IN:S17N (the population of IN:M50I with RH:N79S and IN:S17N were 3.8 % and 1.1 % of the total sequences, respectively). Thus, additional compensatory mutations for IN:M50I may be present in the remaining 149 variants. This population analysis also revealed a number of changes at IN:M50. In additon to IN:M50I, we observed IN:M50L, IN:M50R, IN:M50T and IN:M50V mutations. To define the role of these mutations in viral fitness, variants containing each mutation (HIV(IN:M50V), HIV(IN:M50R), HIV(IN:M50L), and HIV(IN:M50T)) were constructed and replication fitness was assessed. Unlike M50I, those mutants replicated at the same level as HIV(WT) (Supplementary Fig. S5), which highlights the uniqueness of the M50I variant.

### Impact of IN domains on the inhibition of autoprocessing

IN is composed of three domains, the N-Terminal Domain (NTD), a Catalytic Core Domain (CCD), and the C-Terminal Domain (CTD) (32, 33) (Fig. 7a). Codon 50 is located at a linker region between the NTD and the CCD. It has been reported that the CTD of IN regulates assembly and autoprocessing in HIV_NL4.3_ or HIV_HXB2_ strains (34–36). To define the role of the C-terminal domain of IN in the IN:M50I mutant, a series of truncated variants of the C-terminal domain was constructed (Fig. 7a) and GagPol processing was analyzed by WB using purified viral particles using anti-PR and anti-CA. All truncated mutants contained the mature form of PR and CA in the virions (Fig. 7b and 7c). Interestingly, a variant containing an Asn-to-Gly substitution at the C-terminal codon 288 (D288G) rescued the impaired M50I GagPol processing and viral replication (Fig. 7d, Supplementary Fig. S6), and amounts of released virus from the producing cells were comparable to that of wild-type (Supplementary Table S2) indicating that D288 is a critical residue in GagPol homodimerization and autoprocessing in the context of the GagPol(IN:M50I) mutation.

**Figure 7.**
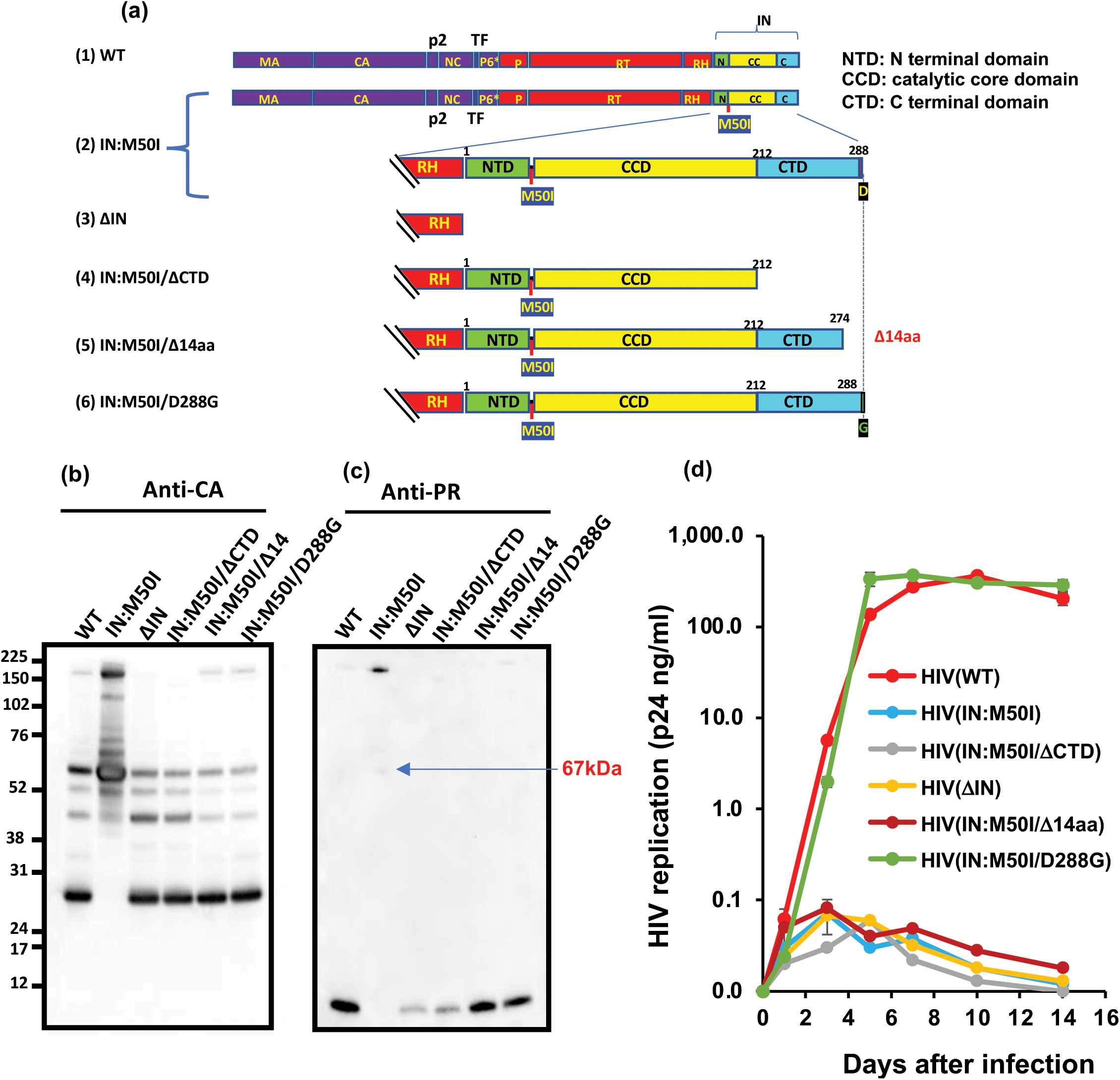
Effect of C-terminal region of IN on HIV(IN:M50I) replication. (A) Schematic structure of HIV mutants. Full-length IN (1–288 aa) was modified using site-directed mutagenesis. HIV(WT) lacking full length IN was designated HIV(ΔIN), HIV(IN:M50) constructs missing either the full-length of the CDT domain and the last 14 aa sequences are designated HIV(IN:M50I/ΔCTD) and HIV(M50I/Δ14aa), respectively. HIV(IN:M50I) with IN:D288G mutation was denoted HIV(IN:M50I/D288G). (b and c) a total viral lysate (5 μg of viral protein) from HIV(WT), HIV(IN:M50I), HIV(ΔIN), HIV(IN:M50I/(ΔCTD), HIV(M50I/(Δ14aa), and HIV(IN:M50I/D288G) were underwent WB using anti-CA (B) or anti-PR (C) antibodies. One of representative results from three independent assays are shown. (D) PHA-stimulated primary CD4(+) T cells were infected with HIV(WT), HIV(IN:M50I), HIV(IN:M50I/DCTD), HIV(DIN), HIV(IN:M50I/ Δ14aa), or HIV(M50I/D288G) and cultured for 14 days with new media every 3-4 days. HIV replication was monitored using a p24 antigen capture assay. Data are representative of three independent experiments with mean ± SD (*n*=3).

### Prediction of Structure Analysis

Mechanims of the assembly of Gag polyporitein are well-investigated (13, 37–41), while a role of GagPol polyprotein is poorly understand. We assumend that IN:M50I mutation may alter the structure of the C terminus region of of GagPol polyprotein, and the compensatory mutations may influence the structure change. To address the hypothesis, we attempted to perform *in silico* Structure analysis. As a full-length of GagPol polyprotein was not crystalized yet, we built a fusion proteins of RT, RH and IN (RT/RH-IN) using the currently available protein prediction programs as described in the Materials and Methods, and then structure analysis was conducted. A total of seven mutant fusion proteins containing RH:N79S, IN:S17N or IN:D288G mutation with or without IN:M50I were archtected, however, none of mutation indicated the significant changes in the models (Supplementary Fig. S7a – S7d).

### Impact of IN:M50I on Alix and Tsg101 incorporation

The mechanisms of virion buddings and virus release have been extensively investigated using Gag polyproteins. HIV hijacks the endosomal sorting complex required for transport (ESCRT)-I, ESCRT-II, and ESCRT-III system in the host cells to facilitate the leasing of virus particles from the infected cells (42–48) (Supplemental Fig. S8a). The p6 domain of Gag polyprotein binds to ESCRT-I, ALG2-interacting protein X (Alix) and Tumor susceptibility gene 101 (Tsg101) in an initiation step of the budding, followed by recruiting ESCRT-III, and then Alix and Tsg101 are incorporated in nascent virus particles (37, 49–53). As shown above, in the presence of IN:M50I mutation, virus release was significantly inhibited and TEM and FRET assays demonstrated that the mutation suppressed the formation of neck/stalk and increased the interaction efficiency of GagPol to Gag (Fig 4e and 4g), indicating that GagPol(IN:M50I) may interrupt the binding of p6 to Alix or Tsg101, or suppress the recruitment of ESCRTs and then subsequently induce the abnormal buds. Thus, it was speculated that the released HIV(IN:M50I) particles might contain less Alix and Tsg101 than HIV(WT). To address the possibility, we performed WB analysis using lysate of HIV(WT) and HIV(IN:M50I) particles and compared the expression amount of each protein. Surprisingly, the incorporation of Alix and Tsg101 were increased in HIV(IN:M50I), compared to that in HIV(WT) (Fig 8), implicating that IN in GagPol may regulate the interaction between p6 and those proteins.

**Figure 8.**
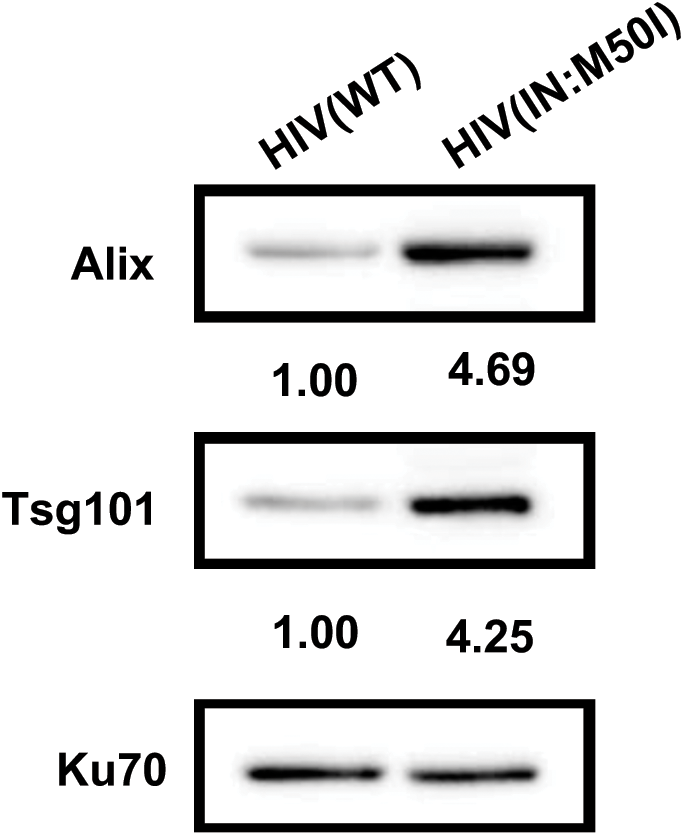
Comparison of Alix and Tsg101 expression in virus particle. A total viral lysate from HIV(WT) and HIV(IN:M50I) were underwent WB using anti-Alix and anti-Tsg101 antibodies. As an internal control anti-Ku70 antibody was used. The intensity of the band for Alix and Tsg101 was normalized by the intensity of Ku70, and the value for Alix/Ku70 and Tsg101/Ku70 are indicated in the image, which is analyzed by Fiji–ImageJ.

## Discussion

Recently, a total of 14 non-synonymous SNPs correlating with different levels of viremia in treatment-naïve HIV-infected patients were reported (11). However, the specific impact of each aa substitution on viral fitness was not investigated. In the present study, we generated HIV-1 variants carrying each of the 14 SNPs to examine the impact of these mutations on viral fitness. We found that the IN:M50I mutation is associated with the loss of replicative capacity. This defect appears to be due to the inhibition of PR autoprocessing by the embedded PR in GagPol polyprotein. Furthermore, we demonstrated that the loss of replicative capacity is rescued by compensatory mutations in either IN or RH, and that the C-terminal residue (IN:D288) contributes to the defect in the IN:M50I context.

It has been reported that GagPol processing is sensitive to the dynamics of viral assembly, and that PR activation is required for correct polyprotein processing (54). Interestingly, our TEM images demonstrated abnormal assembly of virions at the cell membrane and WB studies of the released viral particles illustrated HIV(IN:M50I) mutant also lacked mature PR; a series of FRET analyses illustrated that the GagPol(IN:M50I) can form homodimers and heterodimers with higher efficiencies than GagPol(WT). Although we expected that IN:M50I may change a structure of the C terminal region of GagPol, the *in silco* structure analysis failed to demonstrate any significant changes in the structure of RH-IN fusion protein. To precisly understand the impact of IN:M50I on the increase in the efficiency, we need further structure study.

Infected cells produce both Gag and GagPol polyproteins at a ratio of approximately 20:1 by a −1 frameshifting, which in caused by 5 to 10 % frequency during translation of Gag protein from GagPol mRNA (55, 56) and the ratio of Gag to GagPol proteins in virions is important for Gag and GagPol processing and maturation (56). We demonstrated that GagPol(WT) localized in the cytosole, consistent with other report (17). In contrast, GagPol(IN:M50I) distributed throughout cells (the cytosol and the plasma membrane) as does Gag, suggesting that IN plays a key role in the distribution. The fact that the increase in the localization frequency at the plasma membrane may be responsible for enhancing in the heterodimerization efficiency/frequency and inducing the abnormal assembly.

The molecular events leading to viral assembly and release have been extensively investigated using Gag polyproteins (37, 49, 57–59). Although GagPol polyproteins are also involved in the essential events, a role(s) of the protein has been poorly studied. TEM analysis demonstrated that the accumulation of abnormal buds on the plasma membrane with lack of bud heads and necks/stalks in HIV(IN:M50I)-producing cells (Fig. 2b and Supplementary Fig. S2b). These results suggested that the increase in the efficiency of the heterodimerization of GagPol(IN:M50I) to Gag may disrupt the formation of trimer of Gag polyprotein followed by hexamer of Gag Polyprotein on the plasma membrane (13, 60, 61). The p2 (also known as SP1) region in Gag plays a crucial role in the assembly (40, 62, 63), thus the C-terminus IN:M50I of GagPol may affect the p2 function.

The mechanisms of virion buddings and virus release have been extensively investigated using Gag polyproteins. The p6 domain of the C-terminus region of Gag binds to ESCRT-I, Alix and Tsg101 in an initiation step of the budding; thus, it was also speculated that the abnormality in the assembly may interfere the interaction between the p6 and Alix or Tsg101. Interestingly, WB analysis, however, demonstrated that the HIV(IN:M50I) increased incorporation of the proteins, implicating that IN in GagPol polyprotein may regulate not only distribution of the protein but also the mechanism of the recruitment of ESCRTs. During the recruiting ESCRT-III, in the presence of vacuolar protein sorting 4 (Vps4) at the bud, the formation of ESCRT-III spirals leads to the launch of the head of the budding virus, followed by development of the neck/stalk leading to the final release of viral particles from infected cells (64, 65). IN domain may directly or indirectly regulate the steps (Supplementary Fig. S8b). Further molecular study in the spiral formation will shed light on a role of GagPol in assembly and the release of infectious virus.

Using a series of HIV(IN:M50I) variants containing truncated CTD, we found that the C-terminal aa residue D288 plays a key role in the impaired viral fitnesss in HIV(IN:M50I) mutant. When the acidic hydrophilic aa Asp was changed to a non-polar hydrophobic aa Gly (D288G), viral replicative capability was restored. Unlike other compensatory mutations (RH:N79S and IN:S17N), the D288G alters a net charge in IN, thus the mechanism of the compensation by D288G may differ from that by others. A population analysis using the Los Alamos data base demonstrated that one sequence contained IN:M50I/D288G out of a total of 5100 sequenced samples without IN:S17N or RH:N79S mutations (Supplementary Table S5). Thus, this combination of mutations may be selected. Uncharacterized host factor(s) may interact with the C-terminal of IN and regulate assembly, currently we are investigating the mechanism of the regulation.

A role for IN in proteolytic processing has been reported in previous studies; a selected mutations associated with resistance to an integrase inhibitor KF116 regulated proteolytic processing of Gag and GagPol (8, 66), and quinoline-based allosteric integrase inhibitors (ALLINIs) not only suppress integration but also block the formation of mature virions by impairing their maturation (67, 68). Taken together, these results indicate that the structure of the GagPol polyprotein and the activity of autoprocessing by the embedded PR are intimately linked with the structure of IN. In the present study, we demonstrated that in addition to IN domain, RH domain in the GagPol polyprotein is also invlvoed in the regulation of autoprocessing.

In summary, the current study has demonstrated that naturally occurring polymorphism in the IN gene (IN:M50I) disturbs Gag and GagPol assembly leading abnormal virion formations and suppressing autoprocessing in the released virions and compensatory changes in IN or RH reverses these abnormalities. These findings illustrate the multi-functional aspects of several HIV-1 proteins and indicate that the overall structure-function relationships of the intact virion proteins may involve more than Gag. Further study of functional and structure analyses of GagPol polyproteins and the mechanism of Gag and GagPol interaction may provide new insights into the regulation of viral assembly, budding and viral maturation in HIV, which may disclose new targets to develop novel anti-HIV drugs that may be effective against HIV variants, especiallyl, multi-class drug resistant viruses.

## Materials and Methods

Approval for these studies including all sample materials was granted by the National Institute of Allergy and Infectious Diseases Institutional Review Board and participants were informed written consent prior to blood being drawn. All experimental procedures in these studies were approved by the National Cancer Institute at Frederick and Frederick National Laboratory for Cancer Research and performed in accordance with the relevant guidelines and regulations.

### Cells

Peripheral blood mononuclear cells (PBMCs) were isolated from healthy donors’ apheresis packs using lymphocyte separation medium (ICN Biomedical, Aurora, OH, USA) (69). CD4(+) T cells were purified from PBMCs using CD4 MicroBeads (Miltenyi Biotec, Auburn, CA, USA) according to the manufacturer’s instructions. The purity of the cell types was at least 90%, based on flow cytometric analysis. Cell viability was determined using trypan blue (Thermo Fisher, Waltham, MA, USA) exclusion method. HEK293T cells and HeLa cells were obtained from ATCC (ATCC, Manassas, VA, USA) and maintained as previously described (69).

### Site-directed Mutagenesis

Mutations of interest were induced on pNL4.3, a plasmid encoding a full-length of HIV_NL4.3_ (30) (the plasmid was obtained from Dr. M. Martin through the AIDS Research and Reference Reagent Program, National Institute of Allergy and Infectious Diseases, National Institutes of Health) by the QuickChange Lightinig kit (Agilent Technologies, Santa Clara, CA, USA) according to the manufacture protocol. Briefly, an ApaI (New England Biolab (NEB), Ipswich, MA, USA) and EcoRI (NEB) fragment of pNL4.3 was cloned into the pCR2.1 vector (Thermo Fisher). This clone was used as a shuttle vector and served as the backbone for mutagenesis studies. Primers used in the mutagenesis assays are listed in Supplementary Table S1. All mutagenesis was confirmed by Sanger DNA sequencing using the BigDye terminator v3 (Thermo Fisher) and SeqStudio Genetic Analyzer (Thermo Fisher). After confirming the sequence, the intended clones were digested with *Apa*I and *EcoR*I and then the fragments from *Apa*I and *EcoR*I digestion were used to replace with the corresponding fragment of pNL4.3. DNA sequencing was used to ascertain that each clone possessed the intended mutations.

### Recombinant HIV-1 Viruses

Recombinant HIV-1s were prepared by transfection of pNL4.3 following a method previously reported (70). Briefly, 4×10^6^ HEK293T cells or 2×10^6^ HeLa cells seeded in a 100 mm dish were transfected with 10μg of each plasmid purified using the endotoxin-free plasmid isolation kit (Qiagen, Germantown, MD, USA) with TransIT-293 (Mirus, Houston, TX, USA) for HEK293T cells or TransIT-HeLa MONSTER (Mirus) for Hela cells. Culture supernatants were collected at 48 hours after transfection. After centrifugation at 500 xg for 5min, the supernatants were filtered through a 0.45 μm pore size membrane filter (MiliporeSigma, Louis, MO, USA). Viral particles in the filtrate (8 mL) were ultra-centrifugated on 20 % sucrose in HBS [10 mM HEPES (Quality Biochemical Inc, QBI, Gaithersburg, MD, USA)-150 mM NaCl] cushion for 2 hours at 4 °C and then was resuspended in 80 μL of RP10 or PBS (QBI) and stored at −80 °C until use. Viruses were only used after a single thaw.

Concentration of HIV in each stock was determined by a p24 antigen capture kit (PerkinElmer, Waltham, MA, USA).

### HIV Replication Assay

The levels of HIV-1 replication were determined using primary CD4(+) T cells as follows. CD4(+) T cells were stimulated with 5 μg/mL phytohemagglutinin (PHA; MiliporeSigma) in complete RPMI-1640 (Thermo Fisher) supplemented with 10 mM HEPES, 10 % (vol/vol) fetal bovine serum (FBS; Thermo Fisher), and 50 μg/mL gentamicin (Thermo Fisher) (RP10). The PHA-stimulated CD4(+) T cells (10×10^6^ cells) were infected with 10 ng of p24 of HIV in 1 mL for two hours at 37°C and then cultured at 1×10^6^ cells/mL in the completed RPMI-1640 supplemented with 20 units/mL of recombinant IL-2 (MiliporeSigma) (RP10) for 14 days at 37 °C in T25 flasks. Half of the culture supernatants were exchanged with fresh RP10 every 3 or 4 days of incubation. HIV-1 replication activity was determined by measuring p24 antigen levels in the culture supernatants using the p24 antigen capture assay as described above.

### Western Blot

Virus lysates for Western Blot analysis were prepared using radioimmuno-precipitation assay buffer (RIPA) lysis buffer (Boston biology, Boston, MA, USA) with a proteinase inhibitor cocktail (MiliporeSigma). Total protein concentration in the samples were quantified using a BCA protein assay kit (Thermo Fisher) and Western blot analyses were conducted using the ECL Prime Western Blot Detection system (Thermo Fisher) as previously described (69). Anti-HIV-1 CA antibody (Cat#: ab9071), anti-HIV PR antibody (Cat#:ab211627), anti-HIV1 RT antibody (Cat#: ab63911), anti-HIV IN antibody (Cat#: ab66645) and anti-HIV p55+p24+p17 antibody (Cat #: ab63917) and anti-Tsg101 (Cat#: 30871) were obtained from Abcam (Cambridge, MA, USA), anti-Alix antibody (Cat#: PA5-52873) and anti-Ku70 antibody (Cat#: 4103) were purchased from Thermo Fisher and Cell Signaling Technology (Danvers, MA, USA), respectively. Anti-Nef antibody was kindly provided by Dr. R. Swanstrom through the NIH AIDS Reagent Program, Division of AIDS, NIAID, NIH (Catalog #: 2949) (71). HRP-conjugated anti-Rabbit Ig and anti-mouse Ig antibodies (Cat#:NA931V and NA934V) were obtained from Thermo Fisher.

### Transmission Electronic Microscopic Analysis

Plasmid-transfected HEK293T cells were cultured for 24 hours. Cell-free transection supernatants containing recombinant viruses, or the transfected cells were fixed with 2.5 % Glutaraldehyde (E.M. Sciences, Warrington, PA) in Millonig’s Sodium Phosphate Buffer (Tousimis Research, Rockville, MD) (G-MPB) for 1 min at room temperature. Virus were pelleted as described above and the fixed cells were harvested using a scraper and centrifuged at 500 xg for 15min. The virus and the cell pellets were stored in a fresh G-MPB at 4 °C for overnight. Virus particles were pelleted as described above and then pelleted particle were fixed at 4 °C using G-MPB without disturbing the pellets. All fixed samples were washed repeatedly in Millonig’s Buffer, and then incubated for 2 hours in 1.0 % Osmium Tetroxide (E.M. Sciences), in Millonig’s buffer. Following rinsing steps in Ultrapure Water and en bloc staining with 2.0 % Uranyl Acetate (E.M. Sciences), the samples were dehydrated in a series of graded Ethanol and infiltrated and embedded in Spurr’s plastic resin (E.M. Sciences). Embedded blocks were sectioned using a Leica UC7 Ultramicrotome. 70-80 nm sections were collected on 150 mesh copper grids, and post-stained with Reynold’s Lead Citrate. Samples were examined in a FEI Tecnai Spirit Twin transmission electron microscope, operating at 80 kV.

### RNA Copy Assay

To quantify copy numbers of HIV genomic RNA in virus stocks, the RealTime HIV-I assay kit (Abbot Laboratories, Abbott Park, IL, USA) was used. Briefly, 100 μL of HIV stocks were 10-fold serially diluted and then combined with an internal control RNA from the kit. All assays were conducted on the automated *m2000* System (Abbot) with Abbott mSample Preparation System reagents (Abbot). The detection range of the assay system was 40 to 10×10^6^ copies/mL. BCA protein assay was used to determine total protein concentrations in each sample and the results from the copy assay were normalized by the amounts of protein.

### Förster Resonance Energy Transfer (FRET) Assay

Interactions of Gag-Gag, Gag-GagPol and GagPol-GagPol were measured by FRET assays using a series of the modified-expression plasmids encoding wild type HIV_NL4.3_ Gag or GagPol (lacking frame shifting signal and containing inactive PR with D25N mutation) fused with fluorescent protein genes (EGFP or mSB), generating pGag(MA/EGFP/CA), pGag(MA/mSB/CA), pGagPol(MA/EGFP/CA) and pGagPol(MA/mSB/CA) (17). Since pGagPol constructs lacking the frameshifting signal and *gag* and *pol* were placed in-frame and Gag-Pol polyproteins were translated without the frameshift (17). To induce an inactive PR in pNL(WT) and pNL(IN:M50I), a D25N mutation was induced in the PR of each plasmid using the point mutagenesis with D25N primers (Supplementary Table S1). The pGagPol(MA/EGFP/CA) and pGagPol(MA/mSB/CA) were digested with *ApaI* and *EcoRI* and the *ApaI* -*EcoRI* region was replaced with a corresponding *ApaI* and *EcoRI* fragment of PR-inactive pNL(WT) or pNL(IN:M50I). The subcloned plasmids were termed pGP(WT)GEP, pGP(WT)mSB, pGP(IN:M50I)GFP, and pGP(IN:M50I)mSB. HEK293T cells (50×10^3^ cells) were seeded for 24 hours on μ-Slide 8-Well Glass Bottom chambers (ibidi, GmbH, Planegg, Germany) and then transfected with a total of 0.26 μg of DNA (GFP vector: mSB vector=1:1) using the TransiT-293. 24 hours after transfection, cell images were taken. All images were acquired on a Zeiss Axio Observer.Z1 equipped with the LSM800 confocal module, using a Plan-Apochromat 63x/1.40 objective (Carl Zeiss Microscopy, White Plains, NY, USA); cells were maintained at 37 °C and 5 % CO_2_ during the experiments. For FRET analysis, a combination of three images was taken for each field of view, with respective excitation/emission wavelengths of 488/491-509 nm (Donor (EGFP) channel), 561/587-603 nm (Acceptor (mSB) channel) and 488/587-603 nm (FRET channel). A fourth DAPI image was taken for cell counting purposes, using a 405/410-465nm setup. The pinhole was set at 35 μm throughout all three FRET channels, i.e., 0.82 A.U. for donor channel and 0.70 A.U. for acceptor and FRET channels. Scan zoom was kept at 1.0x resulting in a 0.099 μm pixel size. At least 24 images from three independent experiments per experimental condition were subjected to FRET analysis using the “FRET and Colocalization Analyzer” plugin of the FiJi app, as follows. For each field of view, a bleed-through-corrected FRET index image was generated using the previously described formula (72).

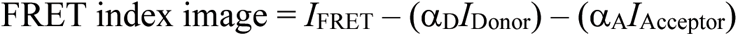

where *α*_D_ and *α*_A_ are signal bleed through coefficients from donor-only and acceptor-only transfection conditions, generated for each independent experiment, and *I*_FRET_ *I*_Donor_ and *I*_Acceptor_ are signal intensities from FRET, Donor and Acceptor channels, respectively; a 1-pixel radius median filter was subsequently applied to resulting FRET index images to smooth signal dispersion. Resulting background levels on filtered images were determined for each independent experiment and filtered-out, thereby creating masks mapping every above-ackground FRET-positive pixel. These masks were then applied to both unfiltered FRET index, Donor and Acceptor images, hence allowing for unbiased analysis of FRET efficiency for each field of view as a whole, and not singled-out cells. Apparent FRET efficiencies were calculated as previously described (17, 73), as a function of acceptor (*E*_A_).

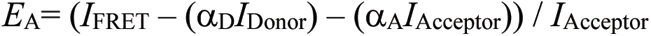

### Tubulin Immunostaining

HEK 293T cells (3×10^5^ cells) were plated on the sterilized round 15mm #1 glass coverslips (VWR) in 12-well plates and seeded for 24hours prior to transfection. Transfection was conducted using 1.5 μg of pGP(WT)GEP, pGP(IWT)GFP or pGP(IN:M50I)GFP as described above and then 24 hours after transfection, culture medium was removed. The cells were fixed with MeOH-free 4% formaldehyde (E.M. Sciences) for 15 minutes at room temperature; fixative was subsequently removed, and cells were washed four times in PBS (Quality Biological). Cells were then permeabilized using 0.1% Triton (Calbiochem, MiliporeSigma) in PBS for 5 minutes at room temperature, then washed four times in PBS, after cells were blocked using the BlockAid™ blocking solution (Thermo Fisher Scientific) for one hour at room temperature, coverslips were then recovered from plates and placed upside-down on a 100mL drop of antibody-containing BlockAid™ (ab204686, Recombinant Alexa Fluor 647 Anti-beta Tubulin antibody at 1:100; ab206627, Recombinant Alexa Fluor 555 Anti-beta Tubulin antibody at 1:100) and incubated in a humidity chamber overnight at 4°C in the dark. Coverslips were then washed 4-times in PBS and mounted on glass slides in a drop of ProLong™ Diamond Antifade Mountant with DAPI (Thermo Fisher Scientific). Imaging was carried out on a Zeiss Axio Observer.Z1 motorized microscope using a Plan-Apochromat 63x/1.40 objective and the LSM800 confocal module.

### Prediction of IN:M50I structure in RT/RH-IN Assembled Structure

To predict structure differences in the presence of IN:M50I, we first constructed *in silico* RT/RH-IN fusion protein. Prior to create the fusion protein, a full-length of homology model of IN structure was architected using the WT HIV_NL4.3_ sequence and the Baker Lab hosted Robetta server following the RosettaCM protocol (74). To construct the homology model of IN based on DNA-free templates, the sequences of three domains of IN, i.e., N-terminal domain (NTD), catalytic core domain (CCD) and C-terminal domain (CTD) (Fig. 7A), were used as query sequences to identify the corresponding templates by BLAST searching the Protein Data Bank (PDB) database (75): Three structures (PDB IDs: 1WJB, 1K6Y and 1WJA) were selected as the templates of NTD with identity of 100 %; three structures (PDB IDs: 2ITG, 1B9D and 1ITG) were identified as the templates of CCD with identity of 99.4 %; and finally two structures (PDB IDs: 5HOT and 1EX4) were chosen as the CTD templates with identity of 100 % and 94.7 %, respectively. These eight templates were then used as the input for Robetta to perform the multiple-template modeling protocol(74). Briefly, the wild type IN (IN:WT) sequence with a length of 288 amino acids (aa) was submitted into the Robetta server and the comparative model protocol was selected with 1000 models sampled. The default options were used for the model building procedure. The structure of the IN(WT) sequence was predicted based on these identified templates where the unaligned regions, such as the very last seven positions in CTD were predicted by using *de novo* fragments. Similarly, to create WT of HIV NL4.3 RNase H (RH:WT) structure, the 120 aa sequence from the NL4.3 genome was submitted into Robetta as the input, with “CM only” option was selected, and 1000 models sampled was used as described above for IN:WT prediction. The multiple templates for RH:WT prediction were automatically identified by Robetta. After having obtained the predicted RH:WT and IN:WT structures by selecting the best model separately, the two structures (domains) were assembled by the AIDA (*ab initio* domain assembly) server (76). AIDA domain assembly procedure consists of several steps: (a) calculate the sequence profile from the full length sequence using PSI-BLAST (77); (b) predict the secondary structure of protein from the profile by PSIPRED; (c) predict solvent accessibility from the output of last two steps; (d) develop the initial full length model in the torsion angle space from the provided domain structures and the predicted secondary structure; (e) optimize the initial model by using knowledge-based statistical potentials where the solvent accessibility derived term plays an important role in guiding the simulation; and (f) select the lowest total energy output from 50 optimized results and add side-chain by SCWRL4 (78) into it as the final model. In the current work, the full-length sequence with 408 aa: 120aa RH:WT and 288 aa IN:WT, and two predicted domain structures (after renumbering the residues as required by AIDA) were submitted into AIDA to predict the two-domain RT/RH:WT-IN:WT structure. After that, seven mutated sequences (RH:N79S, IN:S17N, IN:M50I, IN:D288G, IN:M50I/D288G, RH:N79S and RH:N79S/IN:M50I) and the WT sequence were submitted to Robetta with the same protocol as described above using the predicted RH:WT-IN:WT structure as the single template for modeling corresponding structures.

### Statistical Analysis

Intergroup comparisons were performed using two-tailed unpaired *t*-tests using Prism 8 software (GraphPad, San Diego, CA, USA). *P* values <0.05 were considered statistically significant.

## Acknowledgments

Authors thank Drs. M. Bennedbæk and J.D. Lundgren for providing information of the 14 SNPs to initiate this study, Dr. Y. Morikawa for providing plasmid constructs for FRET assay, Dr. J. Kovacs for discussion, Drs. L. Huzella, X. Jiao, and F. Scrimieri for critical reading of the manuscript. This project has been funded in whole or in part with federal funds from the National Cancer Institute, National Institutes of Health, under Contract No.

HHSN261200800001E. The content of this publication does not necessarily reflect the views or policies of the Department of Health and Human Services, nor does mention of trade names, commercial products, or organizations imply endorsement by the U.S. Government. This research was supported [in part] by the National Institute of Allergy and Infectious Disease. The content of this publication does not necessarily reflect the views or policies of the Department of Health and Human Services, nor does mention of trade names, commercial products, or organizations imply endorsement by the U.S. Government.

## Author contributions

T.I. designed all studies, performed assays, analyzed data, oversaw assays, supervised and wrote the manuscript. Q.C. performed assays. J.G.B., S.L., J.Y., H.H., M.H. and H.S. contributed assays and wrote a draft of manuscript. R.D. and W.C. supervised and reviewed the draft manuscript, H.C.L. conceptualized this project and wrote the manuscript. All authors reviewed the final manuscript.

## Data and materials availability

All data are available in the main text or the supplementary materials.

## Supplementary Information (Figures and Tables)

**Supplementary Fig. S1.**
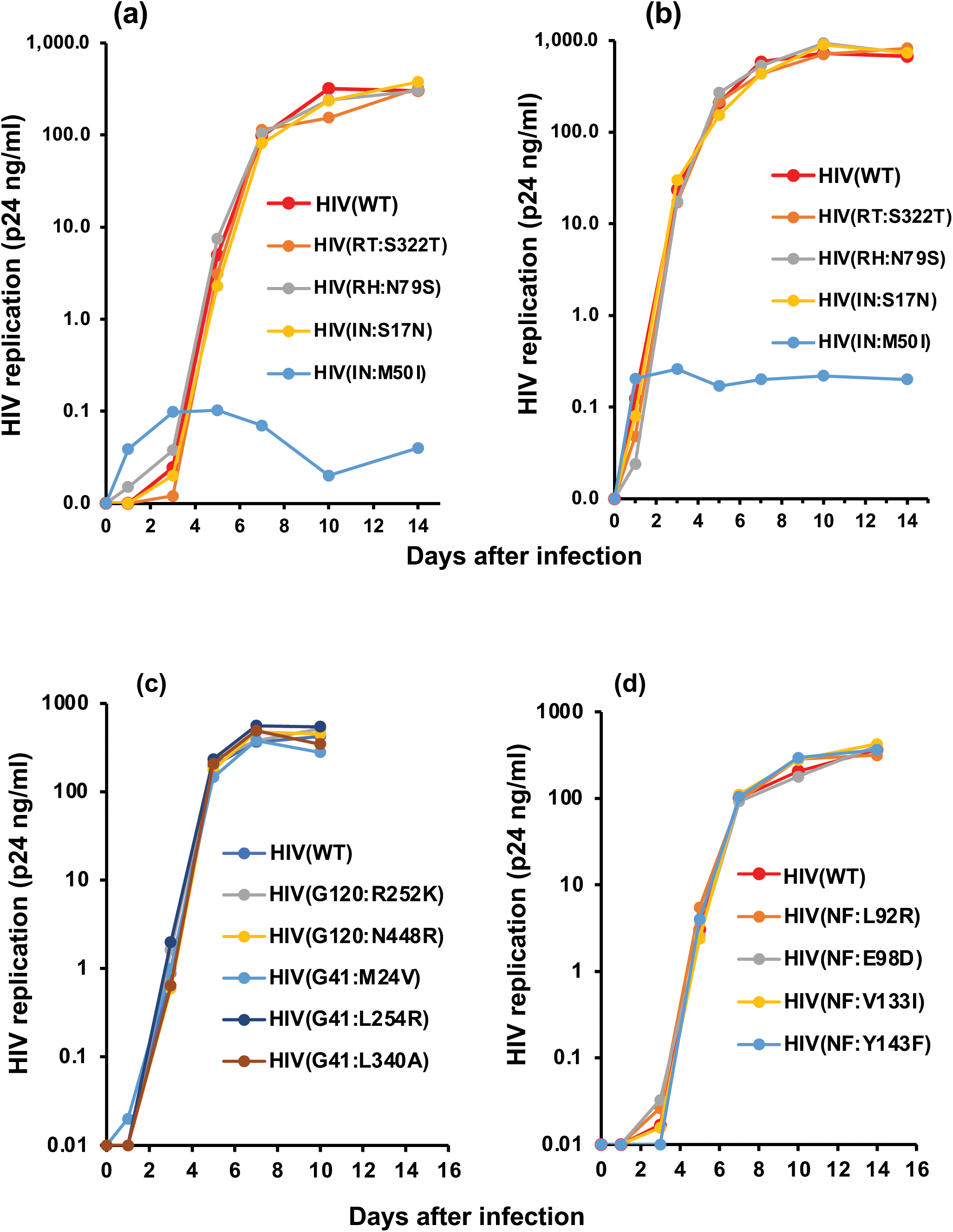
Comparison of HIV replication activity between HIV(WT) and mutants. PHA-stimulated primary CD4 T cells from normal healthy donor were infected with HIV(WT) or variants containing mutation in GagPol as described in the materials and methods. The infected cells were cultured for 14 days with changing media on every 3-4 days. HIV replication was monitored using a p24 antigen capture kit. Data indicates means ± SD.

**Supplemental Fig. S2:**
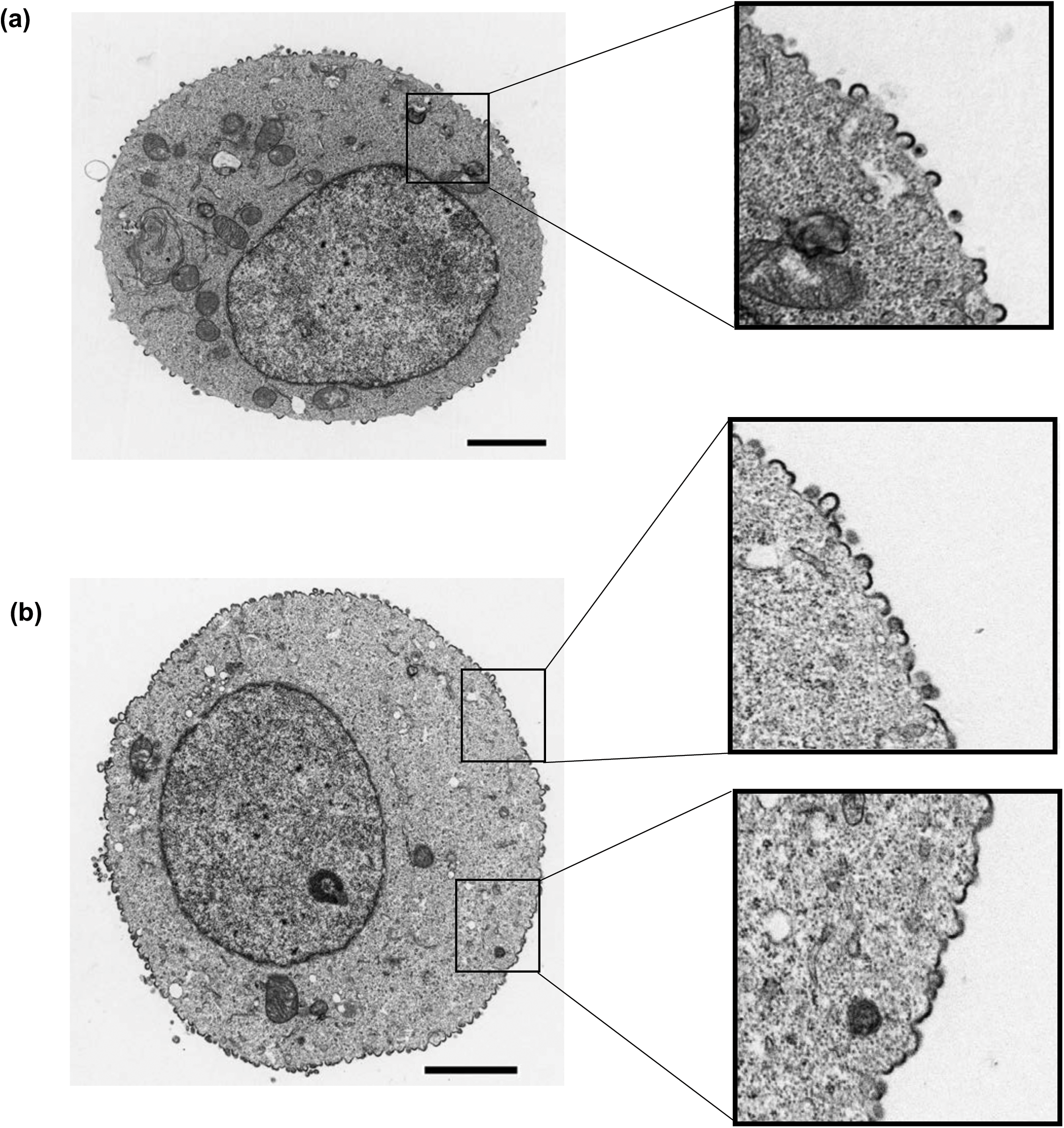
Comparison of cell morphology of HIV(Wt) and HIV(IN:M50I) producing cells. HEK293T cells were transfected with plasmid encoding HIV(WT) (a) or HIV(IN:M50I) (b) gene and cultured for 24 hours each cell was fixed and then subjected for TEM as described in the materials and methods. scale bars indicate 2 µm. Boxes indicate the magnified cell surface.

**Supplementary Fig S3.**
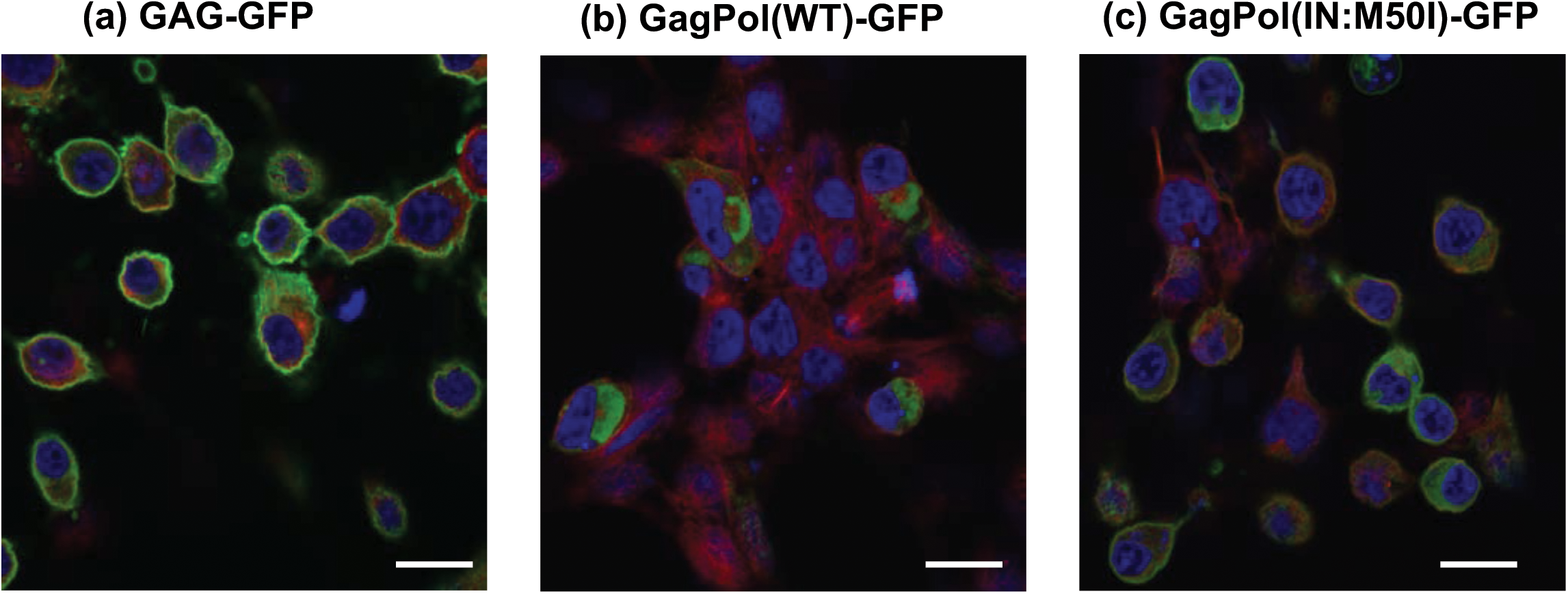
Comparison of Gag and GagPol distribution. HEK293T cells were transfected with pNLGag-GFP, pNLGagPol(WT)-GFP or pNLGagPol(IN:M50I)-GFP construct, and the protein distribution was observed. After transfection, cells were fixed with a fixation buffer (Abcam) and Tubulins were stained with Alexa Fluor® 555 Anti-beta Tubulin antibody [EPR16774] (ab206627). Scale bars indicate 20 µm.

**Supplemental Fig. S4:**
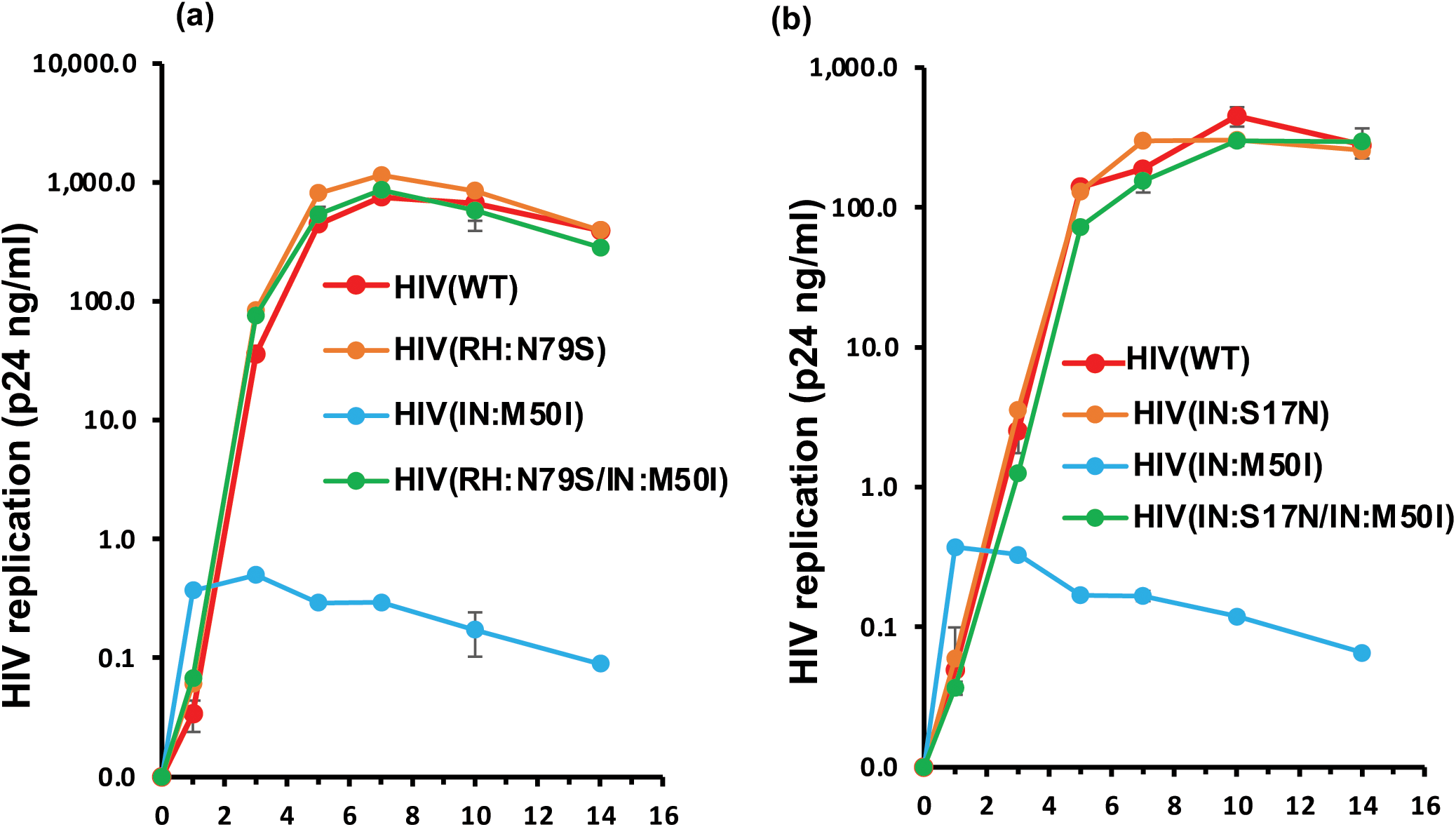
Impact of RH:N79S and IN:N79S on HIV(INM50I) replication. PHA-stimulated primary CD4 T cells from a normal healthy donor were infected with HIV(WT), HIV(RH:N79S), HIV(IN:M50I), HIV(RH:N79S/IN:M50I), HIV(IN:S17N) or HIV(IN:S17N/M50I) as described in the Materials and Methods. The infected cells were cultured for 14 days with changing media on every 3-4 days. HIV replication was monitored using a p24 antigen capture kit. Data indicates means ± SD.

**Supplemental Fig. S5:**
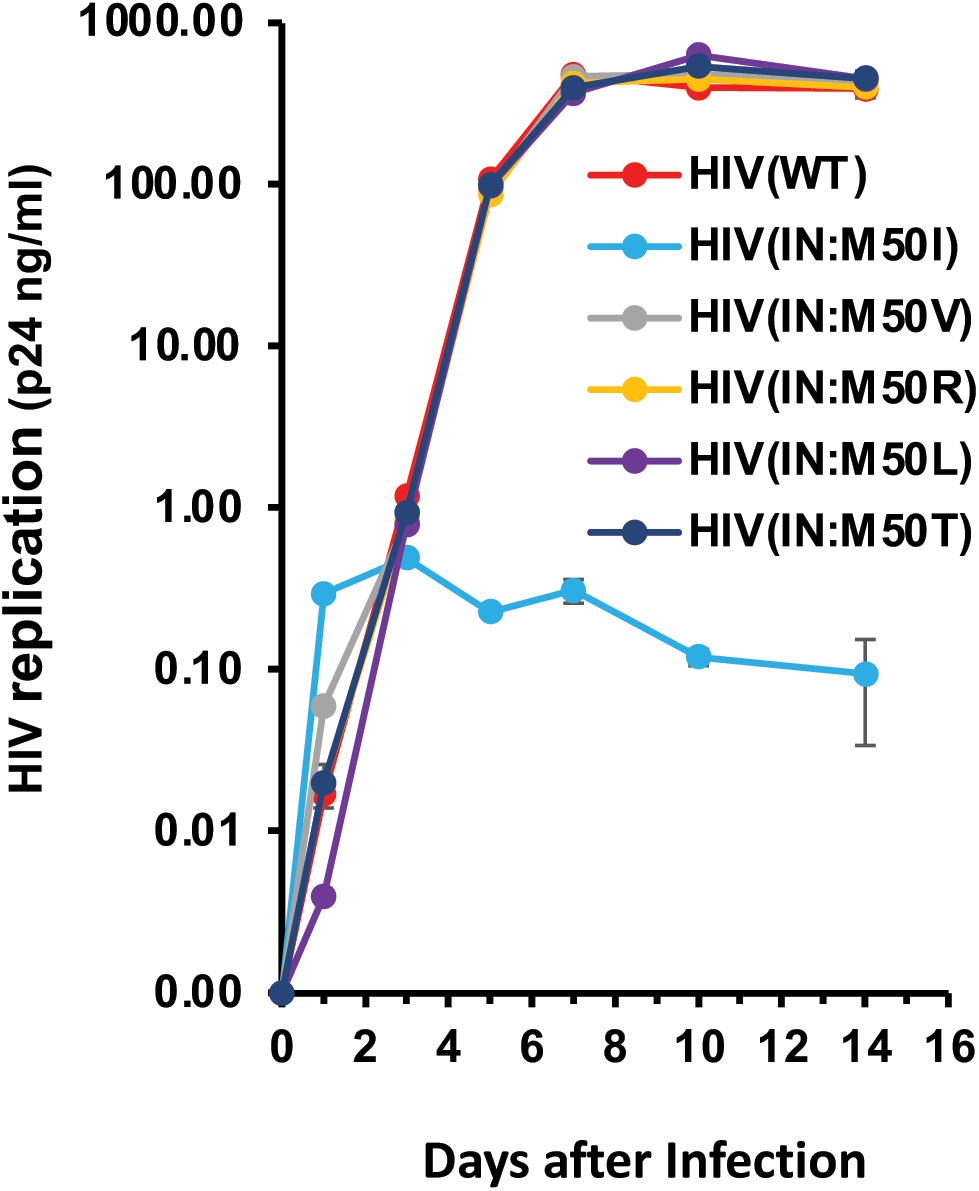
The impact of mutations at codon 50 in IN on HIV replication. PHA-stimulated primary CD4 T cells from a normal healthy donor were infected with HIV(WT) or variants containing a different mutation at codon 50 as described in the Materials and Methods. The infected cells were cultured for 14 days with changing media on every 3-4 days. HIV replication was monitored using a p24 antigen capture kit. Data indicates means ± SD.

**Supplemental Fig. S6:**
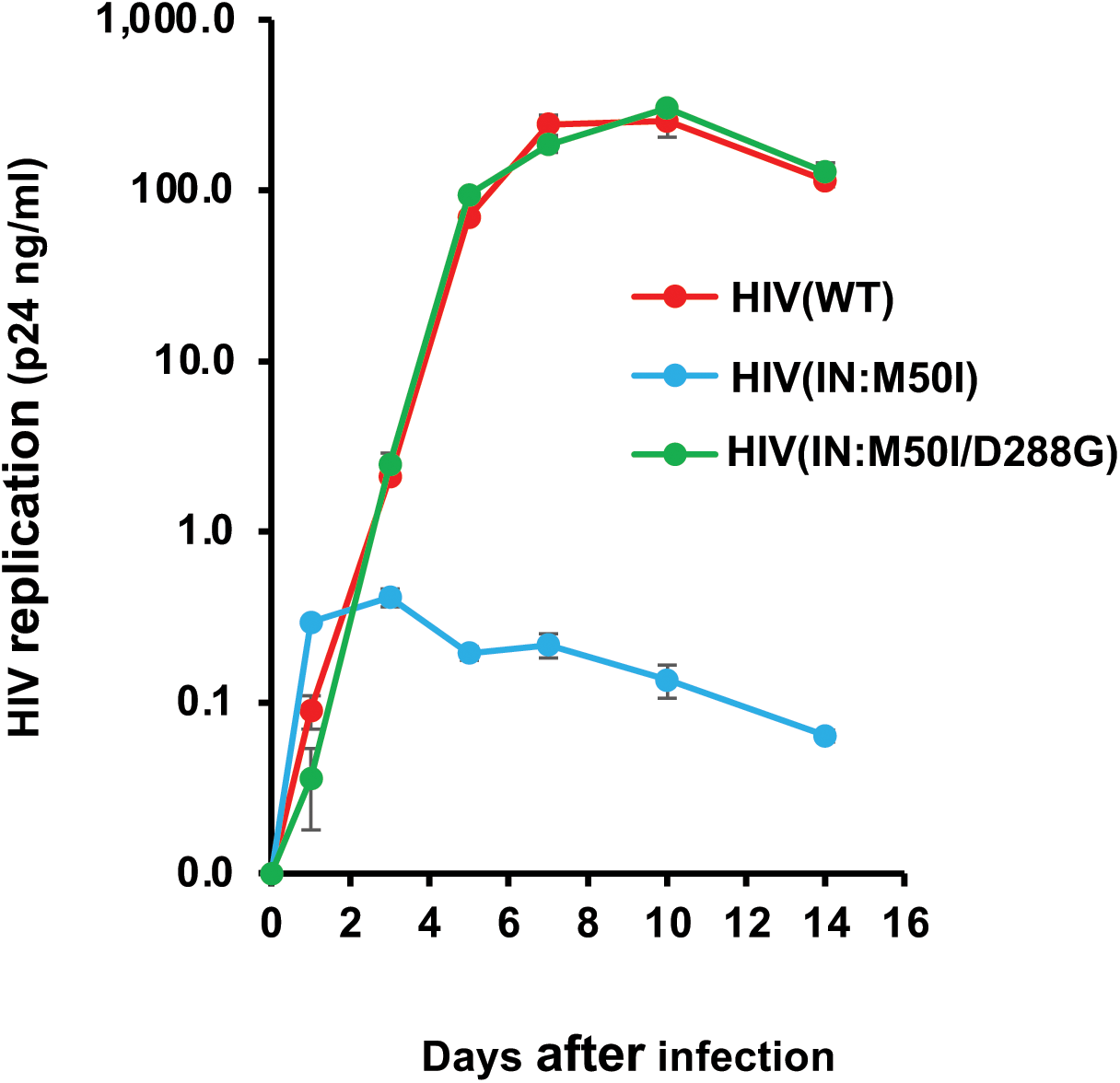
the impact of amino acid change at codon 288 in IN on HIV replication. PHA-stimulated primary CD4 T cells from a normal healthy donor were infected with HIV(WT), HIV(IN:M50I) or HIV(IN:M50I/D288G) as described in the Materials and Methods. The infected cells were cultured for 14 days with changing media on every 3-4 days. HIV replication was monitored using a p24 antigen capture kit. Data indicates means ± SD.

**Supplementary Fig. S7.**
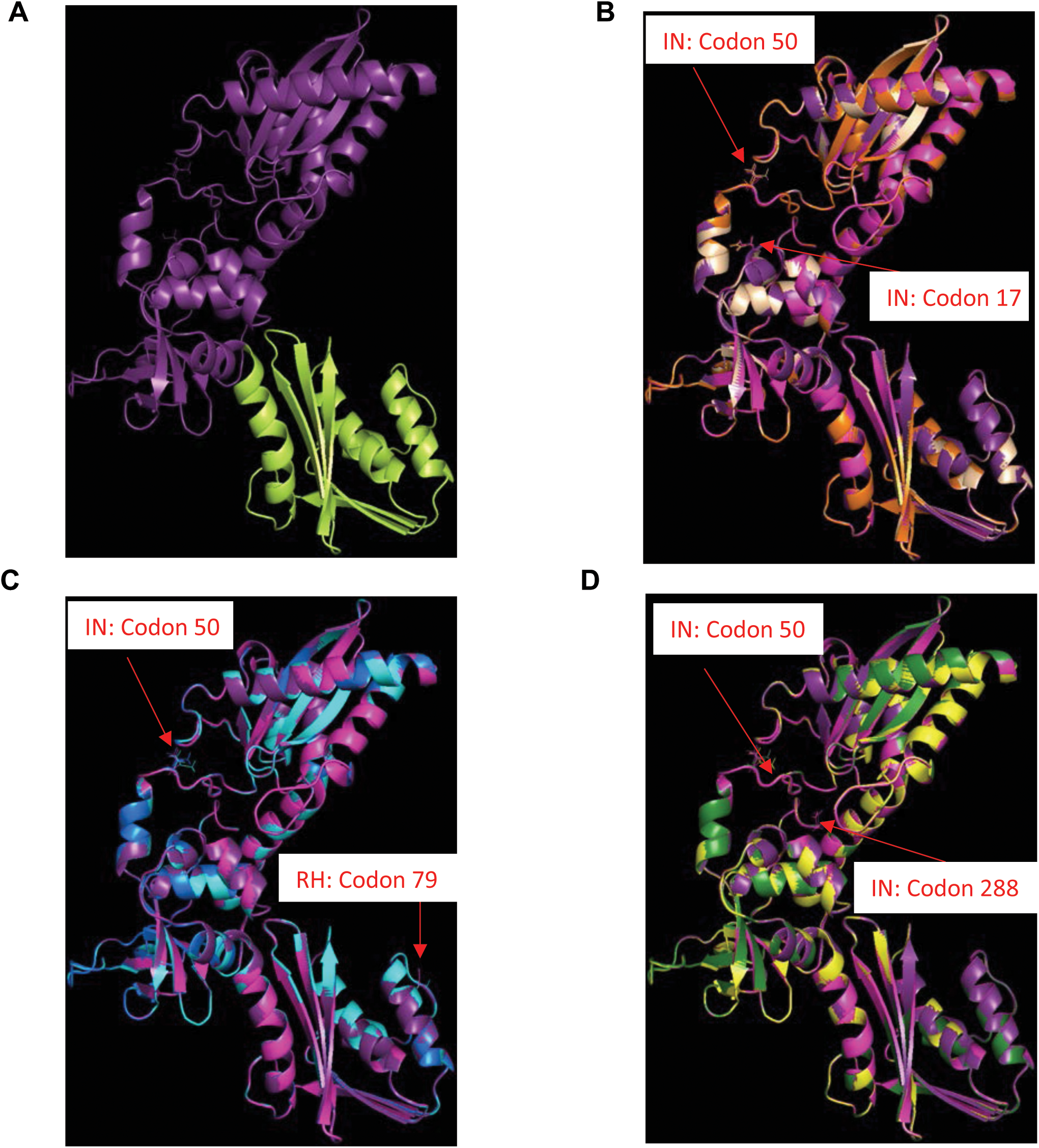
Predicted structure analysis of RH-IN fusion protein. A total of eight *In Silico* RH-IN structures (RH:WT-IN:WT, RH:Wt-IN:S17N, RH:Wt-IN:M50I, RH:WT-IN:S17N/M50I, RH:Wt-IN:D288G, RH:WT-IN:M50I/D288G, RH:N79S-IN:Wt and RH:N79S-IN:M50I) were created and aligned together by using RH:WT-IN:WT as a reference structure. (a) Structure of i*n silico* RH-IN (Wt) fusion protein. IN and RH domains are indicated by is violet purple and lemon, respectively. (b) Alignment of homology models from Robetta among RH:WT-IN:WT (light magenta), RH:WT-IN:S17N (orange), RH:WT-IN:M50I (violet purple) and RH:Wt-IN:S17N/M50I (wheat). (c) Alignment of homology models from Robetta among RH:WT-IN:WT(light magenta), RH:Wt-IN:M50I (violet purple), RH:N79S-IN:WT (cyan) and RH:N79S-IN:M50I (marine). The side chains in positions RH:79 and IN:50 are shown in lines (d). Alignment of homology models from Robetta among RH:WT-IN:WT (light magenta), RH:WT-IN:M50I (violet purple), RH:WT-IN:D288G (yellow) and RH:WT-IN:M50I/D288G (forest). Red arrows show the positions codons.

**Supplemental Fig.S8:**
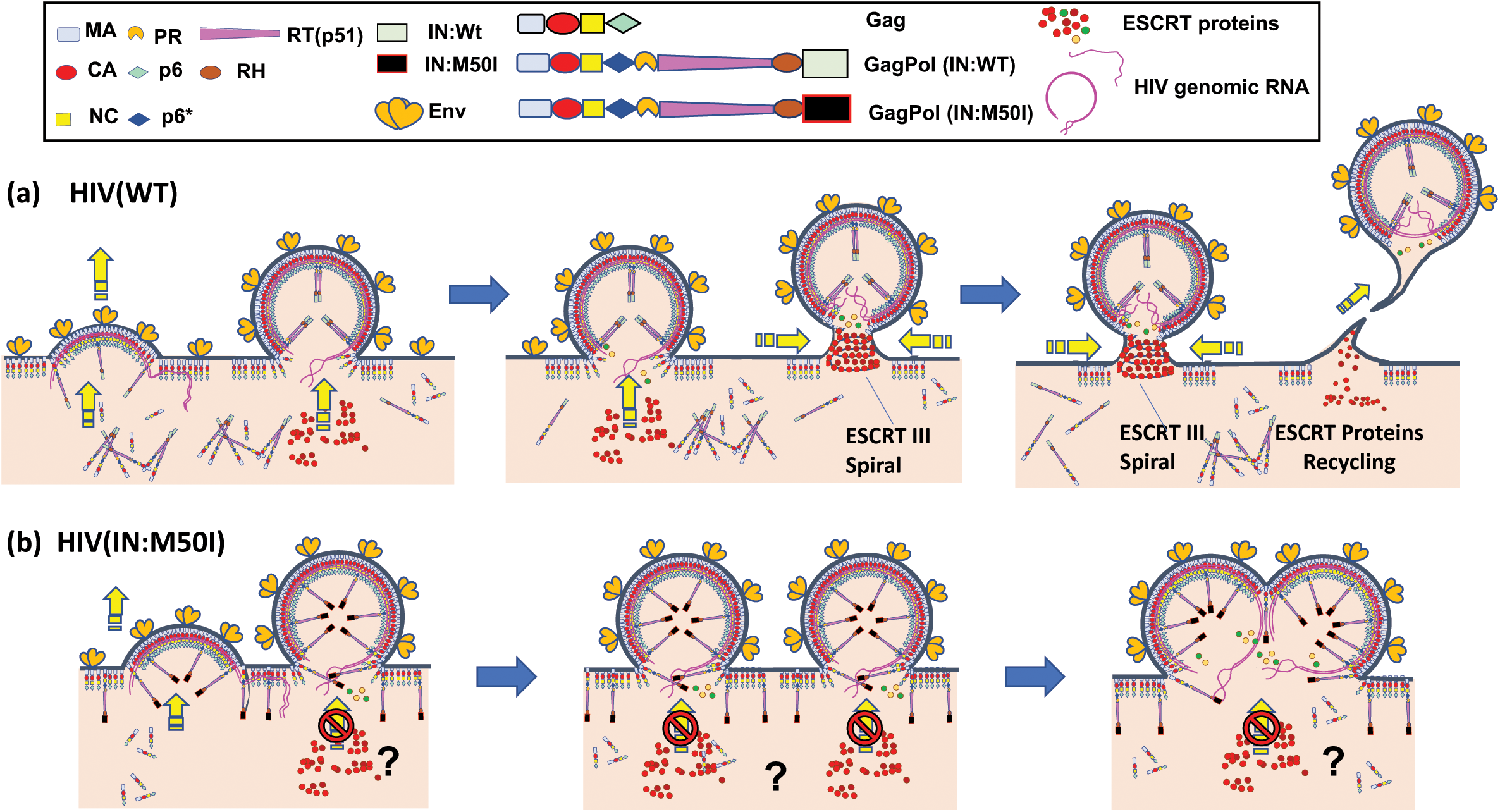
A proposed Mechanism of Suppression of HIV(IN:M50I) Release. In HIV(WT)-producing cells, Gag, GagPol (WT) and viral genomic RNAs(vgRNAs) translocate on the plasma membrane and assemble into membrane-associated lattice, and HIV envelope protein (Env) trimers are incorporated in the budding. Viral release are facilitated by recruiting endosomal sorting complexes required for transport (ESCRT) components. During this process, GagPol forms homodimers. (**b**) In HIV(IN:M50I)-producing cells, GagPol(IN:M50I) efficiently translocates on the plasma membrane and forms heterodimers with Gag, this perturbation of the Gag and GagPol assembly disturbs recruitment of the ESCRT system by uncharacterized mechanism(s) and facilitates to fuse budding particles, subsequently the buds form abnormal size and HIV release is inhibited.

**Supplementary Table 1:**
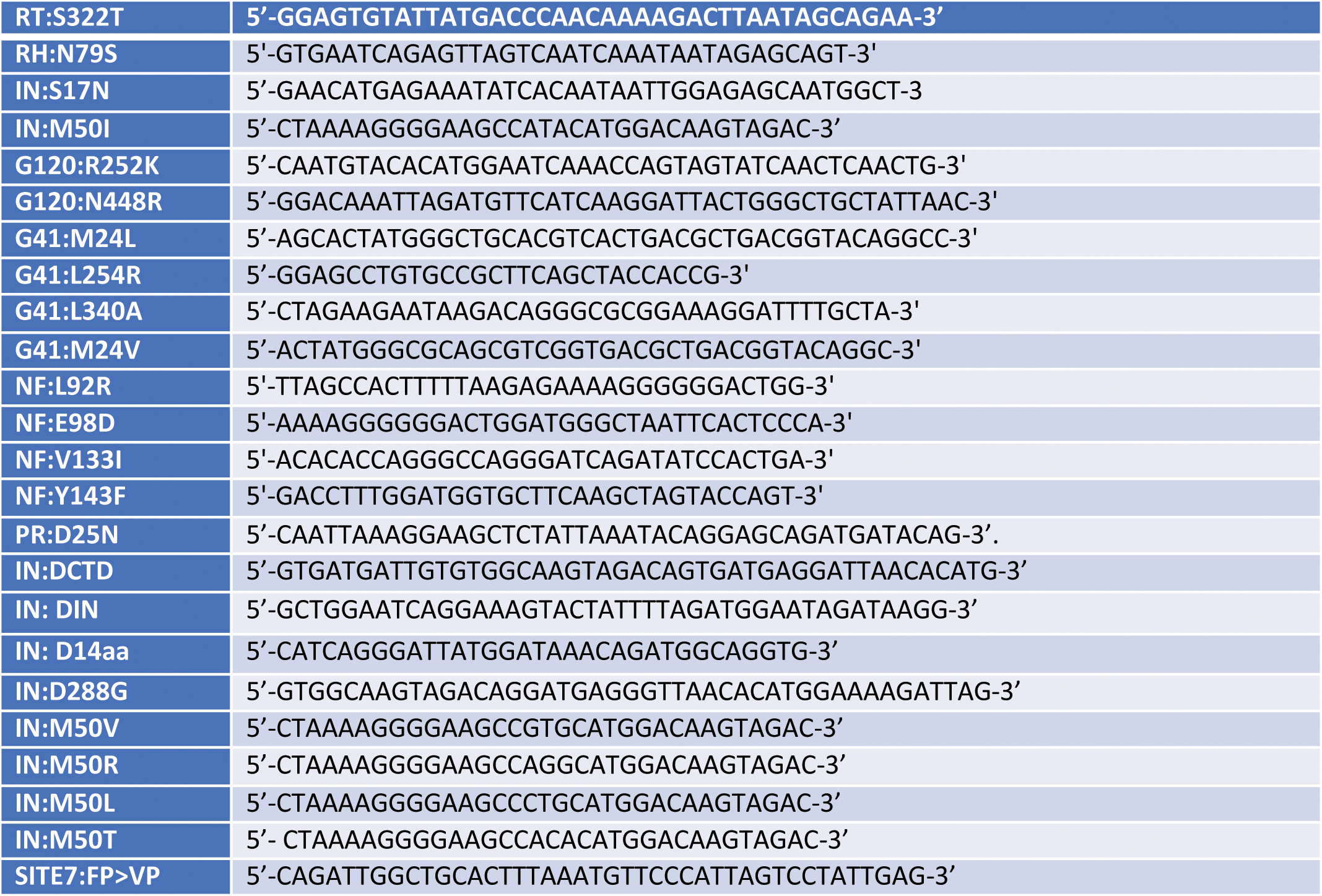
Primer sequence for site-directed mutagenesis (Sense strands)

**Supplemental Table S2:** p24 antigen concentration in HIV stocks.

| Denoted HIV | p24 $\mu\text{g/ml}$ * (%) |
| --- | --- |
| HIV(Wt) | 89 ± 12 (n=7) (100) |
| HIV(RT:S322T) | 71 (n=2) |
| HIV(RH:N79S) | 59 (n=2) |
| HIV(IN:S17N) | 84 ± 14 (n=2) (94) |
| HIV(IN:M50I) | 0.30 ± 0.087 (n=7) (0.3) |
| HIV(G120:R252K) | 79 (n=2) |
| HIV(G120:N448R) | 122 (n=2) |
| HIV(G41:M24V) | 83 (n=2) |
| HIV(G41:M24L) | 85 (n=2) |
| HIV(G41:L254R) | 157 (n=2) |
| HIV(G41:L340A) | 105 (n=2) |
| HIV(NF:K92R) | 128 (n=2) |
| HIV(NF:E98D) | 99 (n=2) |
| HIV(NF:V133I) | 79 (n=2) |
| HIV(NF:Y143F) | 122 (n=2) |
| HIV(IN:S17N/M50I) | 160 (n=2) |
| HIV(RH:N79S/M50I) | 136 ± 28 (n=3) |
| HIV(ΔIN) | 54 ± 8.1 (n=4) |
| HIV(IN:M50I/ΔCTD) | 56 ± 9.7 (n=4) |
| HIV(IN:M50I/Δ14aa) | 124 ± 7.3 (n=4) |
| HIV(IN:M50I/D288G) | 119 ± 22 (n=4) |
| HIV(WT) from HeLa | 296 (n=2) (100) |
| HIV(M50I) from HeLa | 0.825 (n=2) (0.3) |

\* Transfection supernatants from HEK293T cells or HeLa were collected and viral particles were pelleted using ultracentrifugation at 100,000 xg as described in the Materials and Methods.
Each viral pellet was resuspended in 1/100 volume of the transfection supernatants in complete culture media. Viral concentration was determined by measuring HIV p24 antigen by ELISA.

**Supplemental Table S3:**
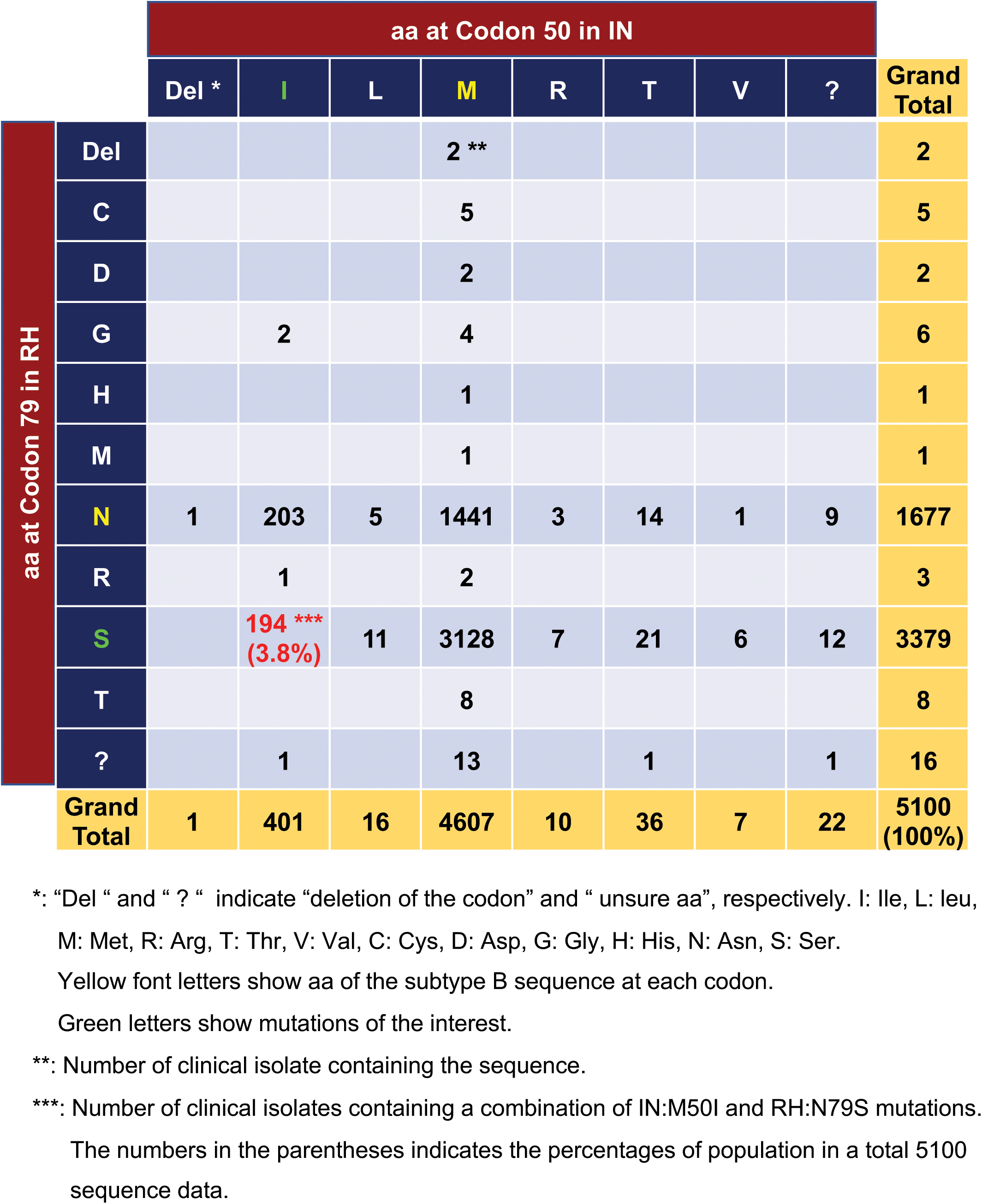
Population analysis of amino acid (aa) residue at Codon 50 in IN and Codon 79 in RH.

**Supplemental Table S4:** Population analysis of amino acid (aa) residues at Codons 17 and 50 and in IN.

|  |  | aa at Codon 50 in IN |  |  |  |  |  |  |  |  |
| --- | --- | --- | --- | --- | --- | --- | --- | --- | --- | --- |
|  |  | Del * | I | L | M | R | T | V | ? | Grand Total |
| aa at Codon 17 in IN | Del |  |  |  | 1 ** |  |  |  |  | 1 |
|  | C |  | 7 |  | 60 | 1 |  |  |  | 68 |
|  | G |  | 1 |  |  |  |  |  |  | 1 |
|  | N | 1 | 58 ***<br>(1.1%) | 4 | 822 |  | 10 |  | 5 | 900 |
|  | R |  | 1 |  |  |  |  |  |  | 1 |
|  | S |  | 323 | 11 | 3607 | 9 | 24 | 6 | 15 | 3995 |
|  | T |  | 9 | 1 | 106 |  | 2 | 1 | 1 | 120 |
|  | ? |  | 2 |  | 11 |  |  |  | 1 | 14 |
| Grand Total |  | 1 | 401 | 16 | 4607 | 10 | 36 | 7 | 22 | 5100<br>(100) |

\*: “Del” and “?” indicate “deletion of the codon” and “unsure aa”, respectively.
I: Ile, L: leu, M: Met, R: Arg, T: Thr, V: Val, C: Cys, G: Gly, N: Asn, R: Arg, S: Ser, T; Thr.
Yellow font letters show aa of the subtype B sequence at each codon.
Green letters show mutations of the interest.
\*\*: Number of clinical isolates containing the sequence.
\*\*\*: Numbers of clinical isolates containing a combination of IN:M50I and IN:S17N mutation.
The numbers in the parentheses indicates the percentages of population in a total 5100 sequence data.

**Supplemental Table S5:**
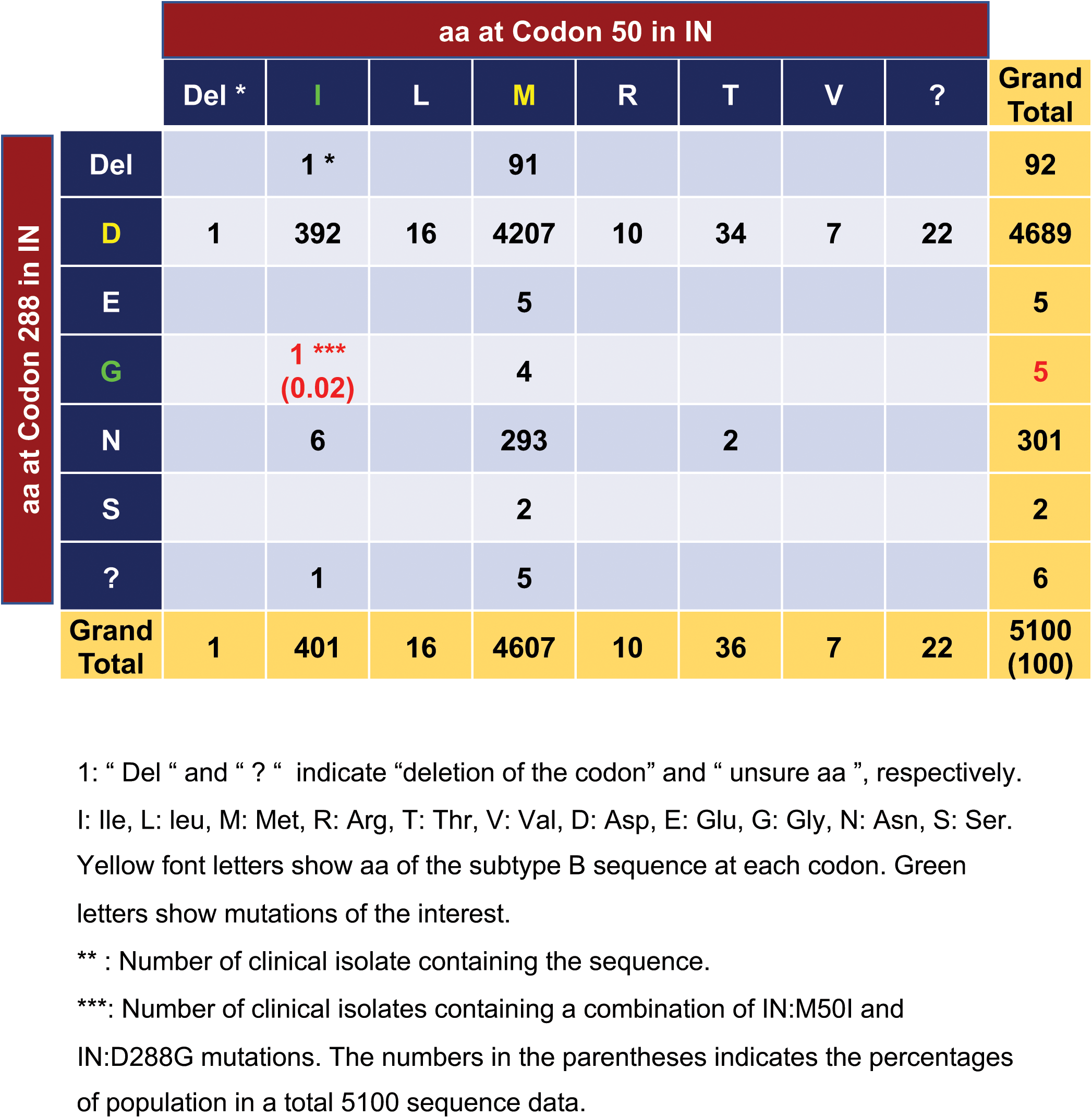
Population analysis of amino acid (aa) residue at Codons 50 and 288 in IN.

